# Triple-N Dataset: Non-human Primate Neural Responses to Natural Scenes

**DOI:** 10.1101/2025.05.06.652408

**Authors:** Yipeng Li, Wei Jin, Jia Yang, Wanru Li, Baoqi Gong, Xieyi Liu, Zhengxin Gong, Kesheng Wang, Zishuo Zhao, Jingqiu Luo, Pinglei Bao

**Affiliations:** Peking-Tsinghua Center for Life Sciences, Peking University, Beijing 100871, China; School of Psychological and Cognitive Sciences, Peking University, Beijing 100871, China; IDG/McGovern Institute for Brain Research, Peking University, Beijing 100871, China; Yuanpei College, Peking University, Beijing 100871, China

**Keywords:** inferotemporal cortex, object recognition, natural scene dataset, neural coding, neural dynamics, cross-species comparison

## Abstract

Understanding the neural mechanisms of visual perception requires data that encompass both large-scale cortical activity and the fine-grained dynamics of individual neurons. While the Natural Scenes Dataset (NSD) has provided substantial insights into visual processing in humans (Allen et al., 2022), its reliance on functional magnetic resonance imaging (fMRI) limits the exploration of individual neuron contributions. To bridge this gap, we present a new dataset: the Triple-N (**N**on-human Primate **N**eural Responses to **N**atural Scenes) dataset, which extends the NSD framework to non-human primates, incorporating single-neuron activity and fMRI recorded from the inferotemporal (IT) cortex. Using Neuropixels probes, we recorded neural responses from five macaques as they passively viewed 1,000 shared NSD images. The dataset includes 59 sessions across 27 sub-regions, capturing over 30,000 visual responsive units. Many recordings were obtained from fMRI-defined category-selective regions, such as face-, body-, scene-, and color-selective areas. Our dataset enables in-depth exploration of neural responses at multiple levels - from population dynamics to single-neuron activity, providing new insights into various aspects of visual processing, including the heterogeneity of object selectivity within functional regions and the temporal dynamics of responses to natural images. Furthermore, the dataset enables joint cross-species analyses, by integrating the macaque neural recordings with human fMRI data, offering a framework for comparing and aligning visual representations across primate species. Overall, our dataset provides a valuable resource for advancing our understanding of visual perception, bridging the gap between large-scale neuroimaging and fine-grained electrophysiological signals, while also facilitating the development of computational models of the high-level visual system.

## Introduction

Large-scale datasets have transformed discovery across disciplines. In artificial intelligence, benchmarks such as ImageNet (Deng et al., 2009) catalyzed the deep learning revolution. In neuroscience, massive neuroimaging initiatives—such as the Human Connectome Project (Van Essen et al., 2013) – integrated structural, functional, and diffusion MRI data from >1,200 participants, enabling systematic maps of brain connectivity and its links to cognition and behavior. These successes illustrate how expansive datasets expose hidden regularities, permit theory-level tests and yield models that generalize.

For visual neuroscience, the Natural Scenes Dataset (NSD) represents a landmark achievement, offering high-resolution functional magnetic resonance imaging (fMRI) data from participants viewing thousands of richly annotated natural images (Allen et al., 2022). NSD has provided critical insights into the functional organization of the visual cortex, supporting investigations into category selectivity (Jain et al., 2023), spatial structure encoding (Sarch et al., 2023), and high-level visual representations (Conwell et al., 2024). The dataset has also enabled progress in brain decoding, facilitating the reconstruction of perceived images and the development of computational models that approximate human visual processing (Khosla et al., 2022; Takagi & Nishimoto, 2022). By bridging cognitive neuroscience and artificial intelligence, NSD drives advances in computational modeling, fMRI data analysis, and neural network architectures inspired by biological vision (Margalit et al., 2024; Prince et al., 2022).

Despite its impact, NSD and similar fMRI-based datasets have inherent limitations. While fMRI provides a valuable large-scale view of cortical activity, it lacks the temporal and spatial resolution necessary to resolve the contributions of individual neurons. Neural computations occur at the millisecond scale within finely structured circuits that cannot be directly inferred from fMRI’s blood-oxygen-level-dependent (BOLD) signals (Badwal et al., 2025; Dubois et al., 2015). However, fMRI uniquely enables precise localization of functional regions, capturing the distributed nature of cortical processing. To bridge this gap, complementary approaches are needed—ones that combine the macroscopic spatial coverage of fMRI with high-resolution neural recordings, while maintaining the use of naturalistic stimuli.

Recent advances in electrophysiological recording technologies now make this possible. In particular, Neuropixels probes have transformed systems neuroscience by enabling high-density recordings of both action potentials (AP) and local field potentials (LFPs) with unprecedented spatial and temporal resolution (Jun et al., 2017; Trautmann et al., 2023). Unlike traditional single or multi-electrode arrays, which sample from a small number of neurons, Neuropixels probes feature hundreds of recording channels on a single shank, enabling simultaneous tracking of thousands of neurons within targeted cortical regions. This capability offers a powerful complement to fMRI: while fMRI provides wide coverage of brain areas, Neuropixels allow researchers to zoom in on the fine-grained dynamics of individual circuits, capturing the richness of population activity underlying perception.

Building on these advances, we introduce the Triple-N Dataset (**N**on-human Primate **N**eural Responses to **N**atural Scenes), an extension of NSD to non-human primates, designed to directly link human fMRI-based studies with large-scale neuronal recordings. Using fMRI-guided Neuropixels probes, we recorded 59 sessions across 27 sites in the inferotemporal (IT) cortex, capturing activity from over 30,000 visually responsive units, including approximately 2,100 well-isolated single neurons. Most of recordings were concentrated in functionally defined, category-selective regions—such as face-selective, body-selective, and color-selective areas—identified via prior fMRI mapping, allowing for targeted investigation of high-level visual processing.

This dataset makes several key contributions. First, it enables an unprecedentedly fine-grained examination of category selectivity in response to natural scenes. Second, it reveals how both intrinsic neuronal properties and stimulus features shape temporal response dynamics, providing a rich resource for studying dynamic visual processing in the IT cortex. Third, by aligning macaque neuronal recordings with human fMRI responses to the same stimuli, it facilitates cross-species comparisons of representational structure and population-level tuning. Finally, it offers a unique platform for benchmarking computational models, revealing differences in how visual and semantic information are encoded across biological systems and artificial architectures. Together, the Triple-N dataset offers a cross-species, multi-scale resource for elucidating high-level visual processing and advancing biologically inspired artificial intelligence.

## Results

### Dense sampling of neuronal responses to natural scenes in IT cortex

We first identified the category-selective areas using fMRI to localize regions selectively responses to face (Fig. 1A, left), body (Fig. 1A, middle), color (Fig. 1A, right), object and scene areas in five macaques. These areas spanned from the posterior to the anterior IT cortex, encompassing over 20 regions in total. Beyond these category-selective areas, we also targeted several regions that did not show significant preference to specific categories, labeled as ‘unknown’ below (see Fig. 1B for all areas registered to a common template). The macaques passively viewed 1,000 natural scene images from the NSD ‘shared1000’ set, along with 24 isolated face (12 macaque and 12 human), body (12 macaque and 12 non-primate animals), and object (12 natural and 12 man-made) images as localizer stimuli to confirm the selectivity of the recorded regions (Fig. 1C). Using fMRI-guided electrophysiological recordings, we captured neuronal responses from over 30,000 units across the five macaques using Neuropixels probes (Fig. 1D; see **Methods**). Juice rewards were given every 1 to 4 seconds to maintain fixation. The images were presented with a 150 ms ON/150 ms OFF trial structure (Fig. 1E). This stimulus duration allowed for 4 - 8 repetitions of each image during a 30 - 40 minutes session, with an average of 6289 trials per session across 59 recording sessions.

**Fig. 1.**
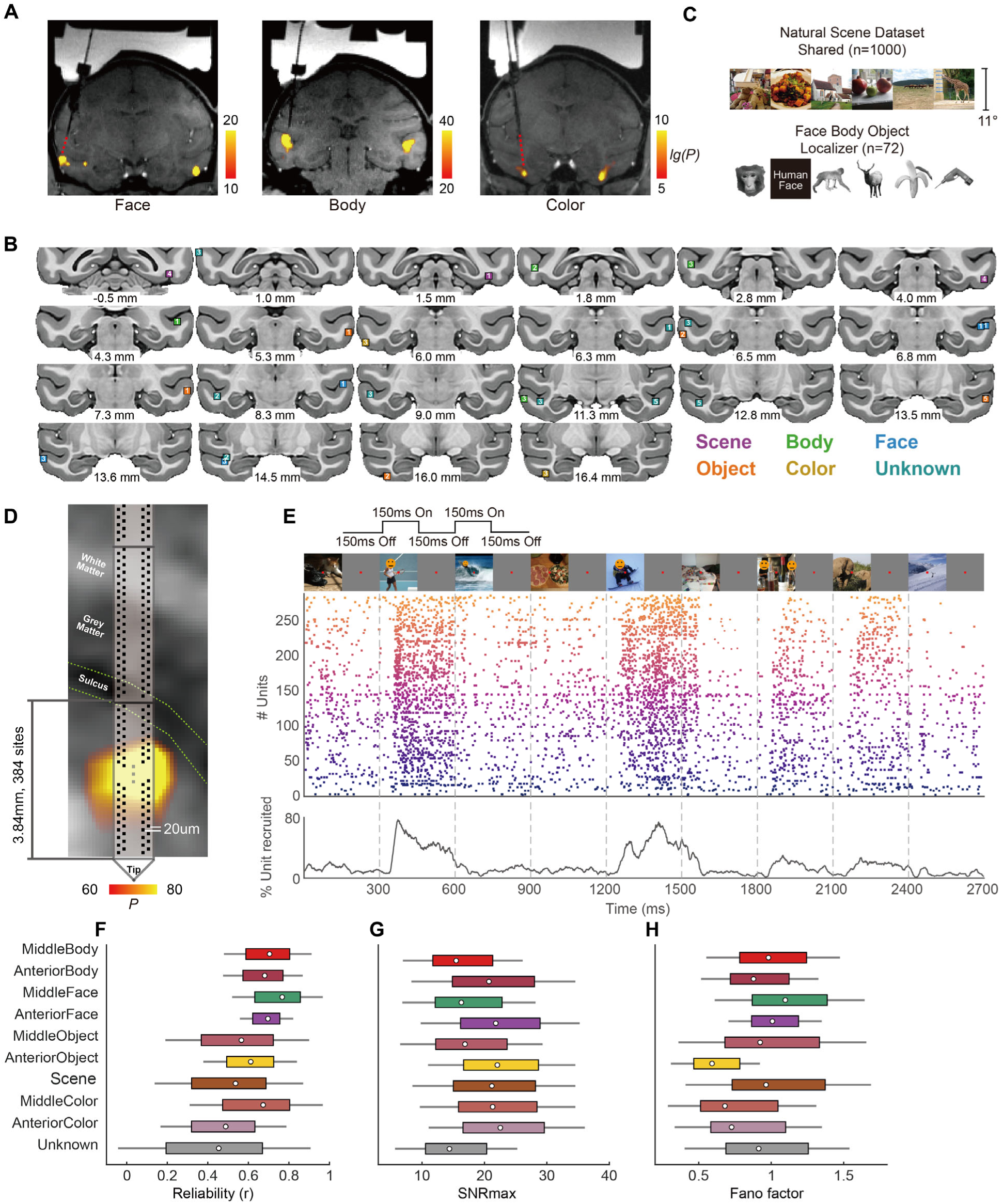
Recording of populational response within categorical-selective regions. **(A)** Coronal slices showing the location of the fMRI-identified areas in M3, targeting the anterior face (left), middle body (middle) and anterior color (right) areas. Red dashed lines indicate expected probe trajectories. Activations for the contrast of specific categories are shown as uncorrected p values. **(B)** Coronal slices showing the targeted brain areas in the NMTv2 template. Numbers below each image represent AP coordinates in the template. Colored squares indicate the recorded areas with color denoting the category selectivity and number within each square indicating the number of subjects. **(C)** Example stimuli used in the electrophysiological experiments. Displayed are six images selected from the 1,000 NSD-shared set and six from the 72 localizer stimuli. **(D)** Illustration of probe placement. Zoomed-in view of the targeting procedure for the middle body area, overlaid on high-resolution anatomical scans. Green dashed lines indicate the boundary between gray matter and the sulcus, serving as key landmarks for guiding probe insertion. **(E)** Example of single-trial populational response. Top, images were presented in a 150ms ON/150ms OFF paradigm. Middle, populational raster plot for 9 example successive trials from middle body area, with unit indices (corresponding to the position of units along the probe shank) coded by color. Bottom, the percentage of neurons recruited during a 30ms time window. Dashed lines indicate stimulus ON time. Due to the bioRxiv policy, all human face images were obscured. **(F-H)** Basic functional properties across regions. ‘Unknown’ refers to all recordings outside functional defined ROIs. In each boxplot, the box marks the interquartile range (IQR), the circle marks the median, and the whiskers represent ±0.5 × IQR. **(F)** Split-half reliability, measured as the correlation between firing rate vectors across halves of the 1,000 image set. **(G)** Maximum signal-to-noise ratio (SNRmax), calculated as the preferred image’s response, minus baseline, divided by the baseline variance. **(H)** Fano factor, calculated as the variance of responses across images divided by the mean response.

We optimized the probe insertion system to ensure stable recordings throughout the experiment, achieving a median probe drift of 6.5 µm across sessions (Fig. S1A, B). Spikes were sorted using Kilosort4 (Pachitariu et al., 2024), and unit quality was automatically assessed with BombCell (Fabre et al., 2023). Units were classified as noise, single units (SU), multiple units (MU), and non-somatic units (non-Somatic, which could be further classified as SU and MU) based on waveform shape, violations of refractory period and spike amplitude (Fig. S1C-G) and other quality metrics. We defined units as visually responsive if they show a significant (*p*<0.001) difference in firing rate between the baseline period (-25 to 30ms relative to stimuli onset) and the response time windows (50–120ms or 120–240ms relative to stimuli onset) using the Wilcoxon rank-sum test. Finally, we sorted more than 30,000 visually responsive units from 59 sessions, including 2,105 SU, 12,406 MU, and 6,510 Non-Somatic units with reliability over 0.4 (for details, see **Methods**). Neuropixels recordings enabled dense sampling of neuronal responses within a ∼4mm distance at 20um vertical spacing (384 channels), yielding simultaneous recordings from hundreds of units and generating reliable trial-wise, populational responses to natural stimuli (Fig. 1E).

To assess data quality, we computed several metrics for each unit in each recording session. First, we calculated split-half reliability (Fig. 1F). Given that response dynamics vary across units (see next section), we employed a moving window approach to identify the optimal period (including start and end times; see **Methods**) for reliability estimation. Cross-validation results indicate that this approach significantly outperforms the fixed time window method (Fig. S1H). We also calculated the max signal-to-noise ratio (SNR_max_) using the most preferred images (Fig. 1H) and the Fano factor for each unit to evaluate image-to-image variability in response to stimuli (Fig. 1G).

With robust measures of response reliability and selectivity, this dataset offers an unprecedented window into neuronal tuning across the IT cortex. While each Neuropixels recording samples only a narrow cortical segment, we overcame this limitation by systematically targeting multiple fMRI-defined patches spanning posterior to anterior IT. Together, this large-scale, spatially distributed recording strategy enables a comprehensive characterization of the fine-scale organization of category-selective responses in the macaque ventral visual pathway.

### Fine-scale structure and variability within category-selective regions

A hallmark of category-selective areas is the high concentration of category-selective neurons (Bao et al., 2020; Popivanov et al., 2014; Tsao et al., 2006). To assess this, we examined category selectivity within these regions using 72 localizer stimuli (Fig. 2A). Given the ∼4 mm recording length of each probe, consistent category selectivity did not always span the entire recorded region. This may be due to the recording trajectory not being perfectly perpendicular to the gray/white matter boundary, causing it to pass through other brain regions rather than being confined to a single category-selective area. To define the category-selective region, we applied a 200-μm spatial bin to the population response and computed category selectivity within each local population. We then defined the category-selective region as the span between the nearest and farthest points exhibiting high selectivity along the recording trajectory (Fig. 2B).

**Fig. 2.**
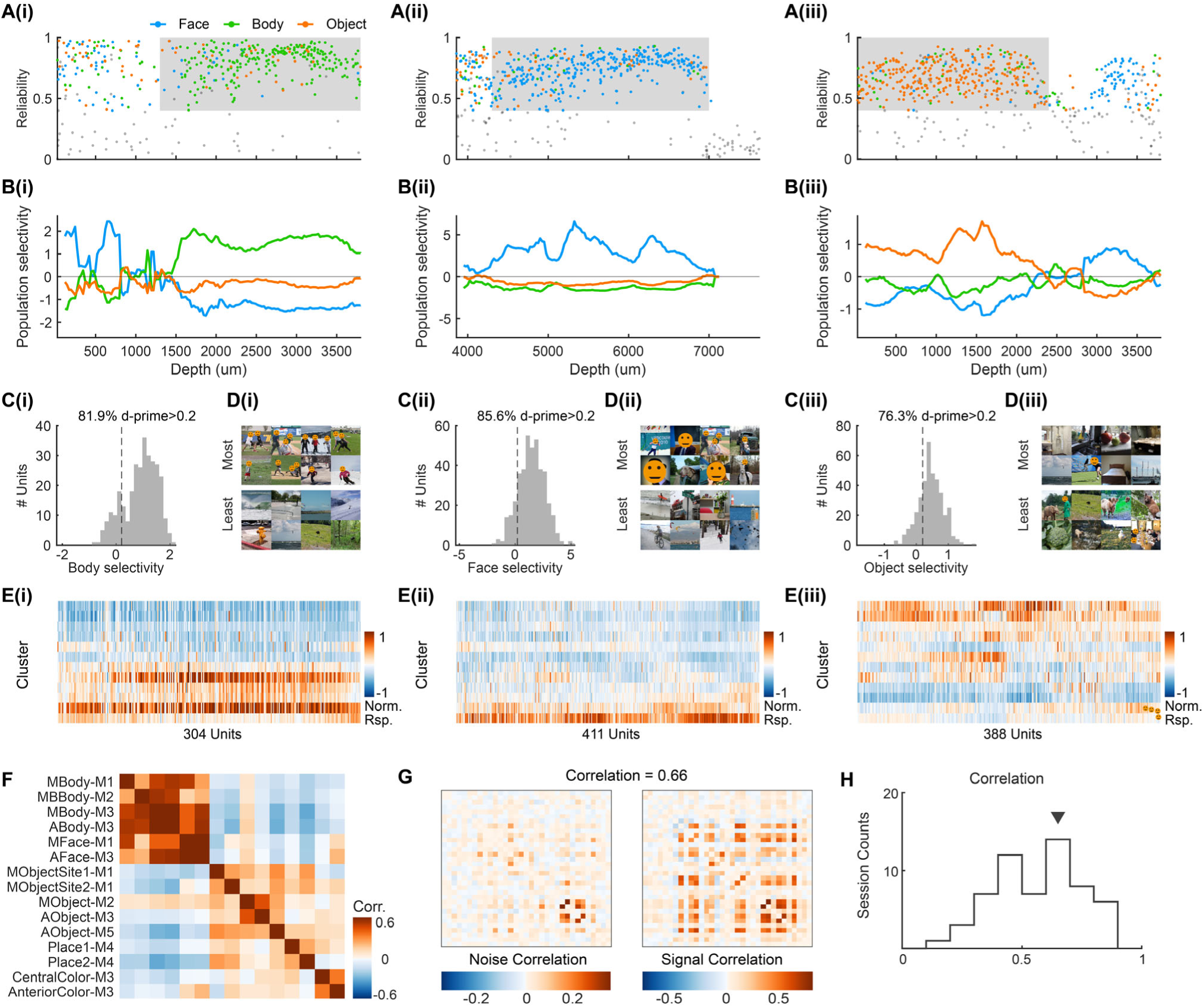
Categorical selectivity profiles across category-selective areas. **(A-E)** Example sessions from body(i), face(ii), and scene(iii)-selective areas. **(A)** Reliability and selectivity as a function of depth, with the grey regions representing identified areas. **(B)** Populational selectivity as a function of depth, where population response was pulled across 200um depth. **(C)** Histogram of categorical selectivity, with a black vertical line representing a selectivity threshold of 0.2. **(D)** Most preferred and most non-preferred images for each area, due to the bioRxiv policy, all face images were obscured with a circle mask. **(E)** Normalized response to each of 12 clusters for each unit, where clusters were ordered by the first principal component of populational response. **(F)** Cross-area similarity was calculated as the Pearson correlation between the mean response profiles of two areas across the 1,000 shared images. **(G)** Example session of pair-wise noise correlation matrix (right) and signal correlation (left)**. (H)** Histogram of spearman correlation values across sessions between noise and signal correlation matrix.

Within these regions, we found that category-selective areas exhibit a high concentration of category-selective units (Fig. 2C). For body and face areas, the proportion is between 74% and 96%. A direct examination of the most preferred images within the population further confirmed categorical selectivity (Fig. 2D). To demonstrate these areas’ category selectivity, we also clustered the shared1000 stimuli based on their semantic labels (see **Methods** and Fig. S2). For each unit, the responses were averaged across each cluster, showing the consistent tuning even for the diverse natural stimuli (Fig. 2E). We also compared the similarities in population-level representations across regions (Fig. 2F), confirming that population preferences align closely within regions sharing the same categorical selectivity, while differing significantly between regions with distinct categorical preferences.

Despite overall consistent tuning within each area, a small but reliable subset of units shows distinct preferences compared to their local populations. As shown in Fig. 2A, most points are marked with the same color, whereas a few points stand out with distinct colors. This observation aligns with previous findings on heterogeneous tuning within category-selective areas (Popivanov et al., 2014). Our dense sampling approach eliminates the need for unit selection, unlike traditional single-unit recordings, thereby enabling a systematic benchmark for investigating the roles of non-selective and even anti-selective units within categorical regions — particularly in the context of circuit-level organization and functional specialization in the primate high-level visual cortex.

Importantly, large-scale, unbiased recordings not only provide a more comprehensive and impartial estimate of neuronal selectivity, but—perhaps more intriguingly—they also offer the opportunity to examine interactions between neurons within the population. One important metric for such interactions is noise correlation, which captures trial-by-trial co-variability between neurons after factoring out stimulus-driven responses. A traditional approach to measuring noise correlations is to present the same stimulus multiple times and compute the mean spike count correlation across trials. However, due to the large number of images tested in our experiment, the number of repetitions per image was limited. To address this limitation, we adapted the functional connectivity approach commonly used in fMRI studies (Norman-Haignere et al., 2012). Specifically, by removing the mean response for each image, we computed correlations between neurons based on the residual trial-by-trial activity (Fig. 2G, left). Next, we examined whether neurons with similar selectivity exhibit higher noise correlations. To do so, we calculated each neuron’s signal correlation (Fig. 2G, right) and assessed the relationship between the signal correlation matrix and the noise correlation matrix. Our analysis revealed a significant positive correlation between the two measurements, indicating that neurons with greater similarity in image selectivity tend to have higher noise correlations (Fig. 2H). Taken together, our results highlight how the Triple-N dataset—enabled by high-density Neuropixels recordings—provides unprecedented access to large-scale, simultaneously recorded neuronal populations, paving the way for systematic investigations of tuning and interaction structure in the IT cortex.

### Diverse coding dynamics across and within neuron populations

Deciphering the neural mechanisms of visual perception requires a dynamic perspective on how neurons process information over time (Kar et al., 2019; She et al., 2024; Shi et al., 2023). Despite the NSD offering extensive insights into visual processing, the sluggishness of the BOLD signal imposes significant limitations in capturing the intricate temporal dynamics. In this study, we leverage our dataset to explore the temporal patterns underlying naturalistic scene processing, offering a unique window into how visual information unfolds over time.

To investigate the dynamic patterns across different units, we first extracted the average peristimulus time histograms (PSTH) in response to 1,000 stimuli for each unit and applied k-means clustering (Fig. 3A). This analysis identified three distinct types of units (Fig. 3B): Cluster 1 (red) exhibited an initial suppression around 70 ms, followed by a gradual rise that peaks at approximately 250 ms, before slowly declining. This pattern suggests a delayed and sustained response, potentially reflecting integrative processing over time. Cluster 2 (blue) displayed a delayed but prominent peak between 140–160 ms, preceded by a minor initial dip and followed by a gradual decline. The response dynamics suggest that these neurons may contribute to later stages of processing, integrating stimulus information at a slightly slower timescale. Cluster 3 (Green) is characterized by a rapid onset and peaks early at around 75–120 ms, followed by a sharp decline. This fast response profile suggests involvement in fast visual encoding, potentially detecting stimulus features or rapid changes in the visual scene.

**Fig. 3.**
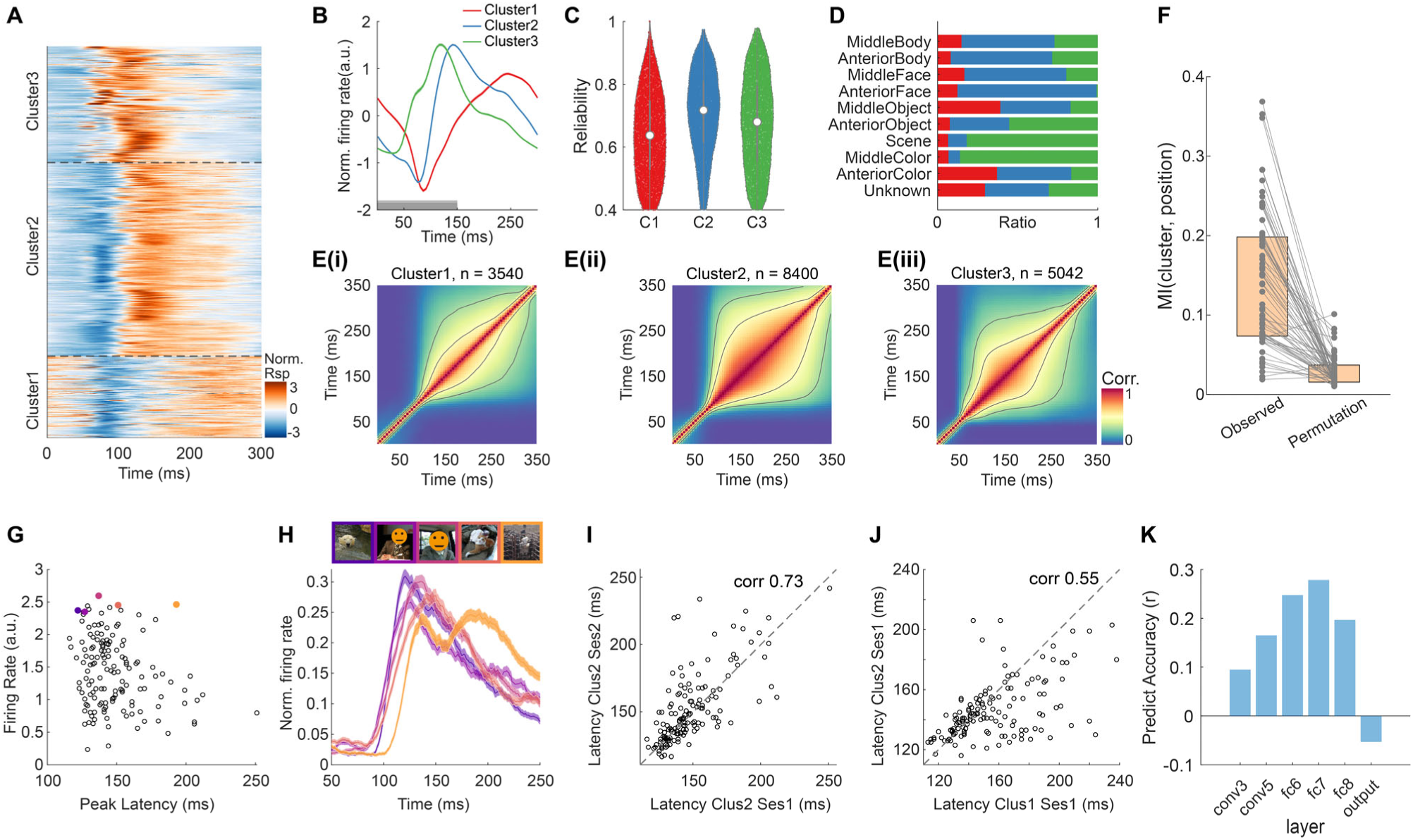
Diverse dynamics of visual processing. **(A)** Normalized firing rate for each unit for one example session. Units were ordered, as defined by cluster type defined with k-means method, with black dashed lines separating different clusters. **(B)** Averaged PSTH for each cluster type. The gray bar indicates the stimulus ON period. **(C)** Violin plots showing the distribution of split-half reliability for neurons in each cluster. **(D)** Distribution of response types across neurons within each recorded region. **(E)** Population-level representation similarity across time. The grey lines indicate contour levels at 0.3 and 0.5. **(F)** Mutual information (MI) between unit type and unit position. Each dot represents a recording session. The box edges correspond to the 25th and 75th percentiles. The left panel shows MI with real data and right panel shows MI with unit position shuffled, *p*<10^-17^, Wilcoxon rank-sum test. **(G)** Population firing rate as a function of peak latency. Colored dots represent images with similar firing rates but different response latencies. **(H)** Five examples of mean PSTHs from a single cluster of neurons recorded in middle face patch. The top panel shows the corresponding images, with color frames matching those in panel **G**. Due to the bioRxiv policy, all face images were obscured with a circle mask. **(I)** Population response latencies from one session plotted against those from another session in the same area, demonstrating the reproducibility of latency patterns across sessions. **(J)** Comparison of population response latencies between two neuron types recorded in the same session. **(K)** Prediction accuracy for peak response latency based on features from different AlexNet layers.

Despite their distinct temporal profiles, all three types demonstrated relatively similar response reliability (Fig. 3C), suggesting that the observed clustering is not merely a consequence of response variability but instead reflects functionally meaningful response dynamics. Moreover, these clusters were widely distributed across all category-selective regions, though their prevalence varied systematically (Fig. 3D). For example, fast-firing units (Cluster 3) were disproportionately concentrated within middle color-selective patches, suggesting a potential functional link between response timing and feature selectivity. The response timing profiles of neurons correspond closely to the representational structures observed across time (Fig. 3E and Fig. S3A), highlighting the reliable encoding of visual information during periods of neuronal activity.

Beyond functional classification, we further examined the spatial organization of these response types and found that their distribution appeared to be linked to their physical positions along the probe shank (Fig. S3B, C). To formally assess this spatial clustering, we computed the mutual information between unit type and physical location and compared it against a permutation-based null distribution (see **Methods**). The results revealed a significantly higher mutual information value (*p* < 10^-17^), indicating that unit types were not randomly distributed but exhibited a structured spatial organization (Fig. 3F). This non-random spatial clustering of units’ temporal profile suggests that neuronal response dynamics may be optimized for local and distributed processing within the visual cortex, potentially facilitating coordinated information flow between units with complementary processing roles. Such an organization could serve to minimize wiring costs, enhance temporal processing efficiency, and support hierarchical integration of visual information within different cortical regions.

The previous analysis focused on identifying distinct temporal response patterns averaged across images. Here, we sought to determine whether different images elicit similar or distinct response profiles within specific unit types. Using Type 2 units from the anterior face area in M3 as an example, we calculated the population-level response timing profile and peak latency for each of the 1,000 images (Fig. 3G). Interestingly, we observed that images with similar overall firing rates could exhibit diverse response latencies (Fig. 3H). A closer examination of the average PSTH across units revealed that responses with the longest latency also exhibited a second-wave response. Notably, the image that evoked this distinct PSTH pattern contained occluded face, suggesting that these stimuli may require recurrent computations or additional visual integration, leading to delayed but sustained neuronal responses. More broadly, we found that these latency characteristics were highly reproducible across recording sessions (Fig. 3I) and across unit types (Fig. 3J), demonstrating their robustness as an intrinsic property of the neural response. Furthermore, these latency properties were shown to be predictable using computational models (see **Methods**, Fig. 3K), indicating that systematic image features contribute to shaping the temporal structure of neuronal responses. These results demonstrate that latency variations are systematic, reproducible, and predictable.

Finally, we observed a relationship between population-level response timing and neuronal preference. Specifically, the initial dip in Type 1 units was more prominent for non-preferred images (Fig. S4), highlighting the potential to link image-specific latency features to underlying visual properties. These findings suggest that different images not only drive variations in firing rate but also shape distinct temporal processing trajectories, which may play a key role in visual recognition.

In summary, our findings reveal that the temporal structure of neuronal responses is shaped by both intrinsic unit properties and stimulus characteristics. This underscores the rich temporal structure embedded in our dataset, offering a valuable resource for investigating how visual information is processed and represented over time in the IT cortex. By capturing fine-grained temporal variations, this dataset provides key insights into the mechanisms underlying dynamic visual processing and serves as a foundation for future computational studies.

### Cross-species comparison between macaque and human high-level visual cortex

Our Triple-N dataset also provides a unique opportunity to compare neural representations with human data collected under the same stimulus. While electrophysiological recordings offer high temporal resolution at the single-unit level, human fMRI captures large-scale population activity across the entire cortex. Despite these methodological differences, both datasets reflect the underlying neural coding of visual information, allowing us to investigate shared and species-specific aspects of visual processing. A particularly compelling domain for such comparison lies in high-level object vision. Macaque IT cortex and human ventral temporal cortex (VTC) were considered as homologous regions due to their similar anatomical locations and their functional specialization in high-level visual processing (Orban et al., 2004), despite existing work has examined this cross-species similarity in organizational (Tsao et al., 2008) and representational (Kriegeskorte et al., 2008) aspect, a direct comparison between these regions’ responses to rich and naturalistic stimuli has yet to be conducted. In this study, we aimed to bridge this gap by comparing the response properties of neurons in the macaque IT cortex with the voxels’ response from the human VTC, using shared 1000 image set.

We first examined the populational-level tuning similarity between species using representation similarity analysis (RSA). We correlated representation dissimilarity matrices (RDM) constructed from BOLD responses from human high-level visual cortex (see **Methods**) and from populational firing rates of neurons in the macaque IT cortex (Fig. 4A). The Spearman correlation between the human VTC and macaque IT RDMs revealed a partial representational similarity between the species (Fig. 4B, left). We further compared this similarity in a dynamic way (Fig. 4B, right) and found that the shared structure remained relatively stable across a broad time window during the trials. However, this inter-species similarity did not fully reach the cross-subject noise ceiling observed within the human data. This discrepancy may reflect systematic differences between the macaque and human visual systems. Future work could aim to identify the specific feature properties that are conserved across species and those that diverge.

**Fig. 4.**
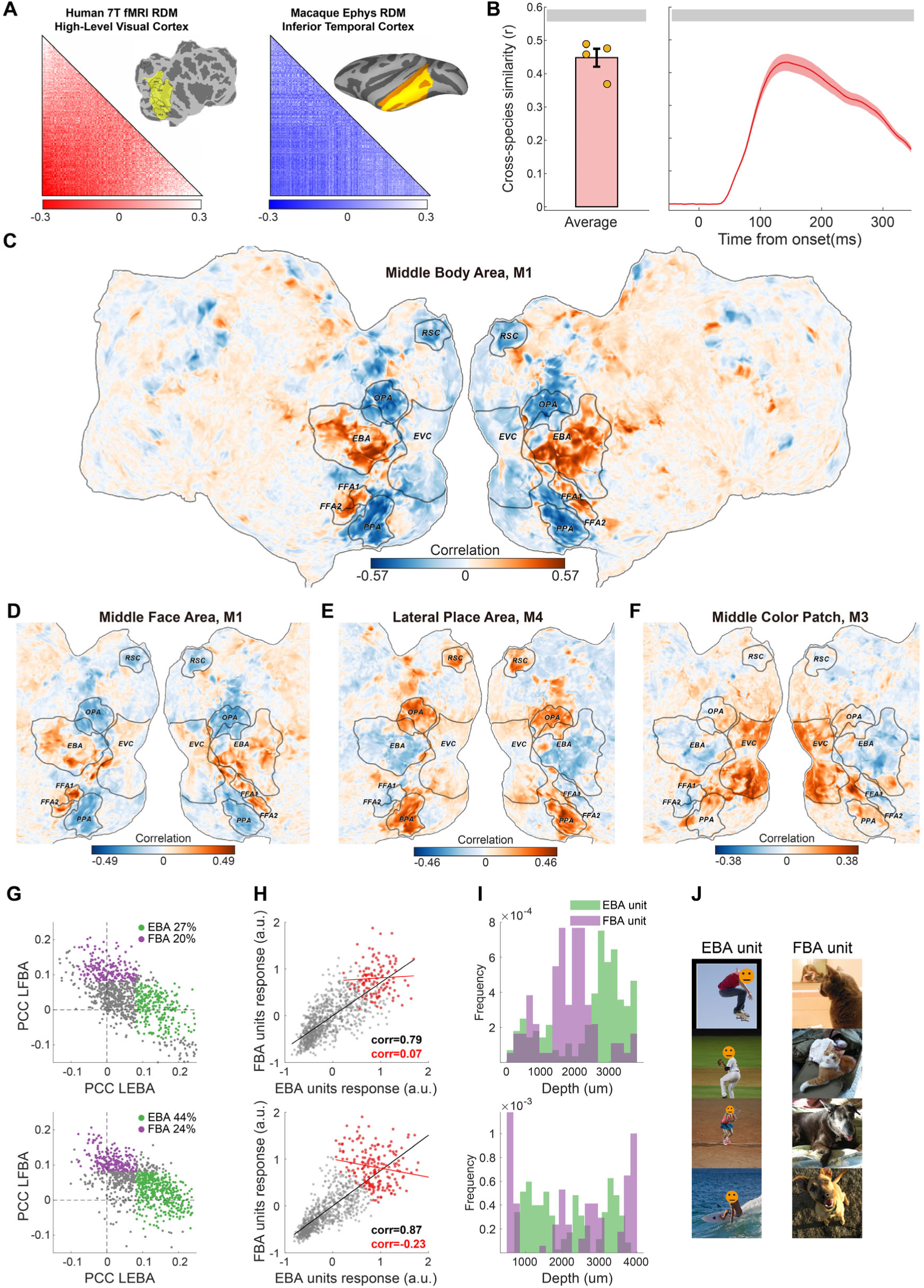
Cross-species comparison of high-level visual cortex. **(A)** Illustration of representation dissimilarity matrices. The left panel shows the mask used for the human high-level visual cortex from one example NSD subject, and the right panel illustrates the macaque IT cortex. **(B)** Representation similarity between macaque and human. Left: Spearman correlation between RDMs constructed from time-averaged macaque electrophysiology responses and human fMRI data, with each dot representing one human subject. Right: Temporal dynamics of Spearman correlation between RDMs computed from macaque neural responses (20ms bins) and the NSD fMRI RDM. The gray bar indicates the cross-subject noise ceiling, and error bars represent the cross-subject SEM. **(C-F)** Neuron-Voxel mapping. Population-averaged responses to 1,000 shared images in macaque category-selective areas were correlated with BOLD responses at each voxel and projected onto the native cortical surface of Human Subject 5. Boundaries of functional regions of interest (ROIs) are outlined in gray. Areas are from: **(C)** the middle body area, **(D)** the middle face area, **(E)** the scene area (LPP) **(F)** the central lateral color area. **(G-J)** Cross-species mapping reveals distinct clusters within the middle (upper panel) and anterior (lower panel) body areas. **(G)** Scatterplot of partial correlations to the EBA and FBA, with colored dots indicating significant (p<0.01) partial Pearson correlation coefficients to specific ROIs. **(H)** Scatterplot of average responses from two neuron types. Gray dots represent all stimuli, while red dots highlight preferred stimuli, defined based on pooled average responses across the population. The gray line shows a linear fit across all stimuli, with a high r value indicating consistent category-level selectivity between the two neuron types. In contrast, the red line fits the responses to preferred stimuli, where a lower r value reflects divergent object-level selectivity between the two groups. **(I)** Spatial distribution of two types of units for an example session. **(J)** Preferred stimuli were defined based on the average responses pooled across the two neuron types. Due to bioRxiv policy, all face images were obscured with a circular mask.

To further examine the correspondence between category-selective regions in macaques and humans, we compared response patterns between macaque firing rates and human BOLD responses. For each category-selective area, we computed the mean evoked firing rates across the 1,000 NSD images in macaques and calculated the Pearson correlation value between these responses and the activation of each vertex in the human cortex. This analysis revealed a strong cross-species correspondence in category-selective representations (Fig. 4C–F). For example, the macaque middle body area showed high correlation with the human extra-striate body area (EBA) and fusiform body area (FBA); the macaque face patch aligned with the human fusiform face area (FFA); and the macaque lateral place patch (LPP) exhibited strong similarity to the human occipital place area (OPA) and para-hippocampal place area (PPA). Additionally, we observed that responses in the macaque central lateral color patch (CLC) were highly correlated with cortical areas between FFA and PPA in humans, consistent with previous findings on human color-selective regions in the ventral temporal cortex (Lafer-Sousa et al., 2016). These findings provide further evidence of a homologous functional organization in high-level visual cortex across primates, suggesting that the primate brain relies on shared computational principles for high-level visual processing.

Beyond the population-level homologous function of category-selective areas, the neuron-to-voxel correlation approach also provides insights into the within-region diversity of local neuronal populations. For example, we examined four body-selective units from a single region and observed distinct correlation patterns. Some units exhibited specialized correlations with the EBA, consistent with population-averaged results (e.g., unit 137 in Fig. S5A), whereas others showed divergent correlation patterns beyond the human VTC, including correlations or anti-correlations with the early visual cortex (e.g., unit 100, 115 and 159 in Fig. S5A). Further analysis confirmed that these units were indeed body-selective but exhibited varying response differences to non-preferred images (Fig. S5B), suggesting that our dataset captures information beyond categorical selectivity. This observation aligns with previous studies macaques face patch AF, which was found to contain diverse functional subpopulations within a single brain area, with some groups showed selective correlation to face patches and some groups showed general correlation with the entire cortex (Park et al., 2017).

To further examine the diversity of IT units, we analyzed the correspondence between single neurons from macaque body patches and multiple body-selective regions in the human brain. We took two body-selective patches from the left hemisphere of monkey M3 served as examples. For comparison, mean fMRI responses to the same 1,000 natural images were extracted from two human body-selective areas— the EBA and FBA—in the left hemisphere. Partial correlation coefficients (PCCs) were then computed between single-unit firing rates and each human region, controlling for variance explained by other brain areas (Fig. 4G). For most units, PCCs with both EBA and FBA were positive, suggesting joint modulation. However, a subset of neurons exhibited selective correspondence, showing a significant PCC with one area but not the other. These patterns remained stable under split-half validation (Fig. S5C), indicating consistent preferences at the single-unit level. This diversity was observed in both middle and anterior body patches: in the middle patch, 27% of units were uniquely explained by EBA and 20% by FBA; in the anterior patch, 44% were uniquely explained by EBA and 24% by FBA. Importantly, this functional diversity also emerged in visual preferences: although population-level preferences for the 1,000 images were strongly correlated between EBA- and FBA-aligned neurons, preferences for the most selective images (defined by average normalized responses > 0.5) were only weakly correlated—or even anti-correlated— between the two populations (Fig. 4H). These functionally distinct neuron types also exhibited spatial clustering within each body patch (Fig. 4I), consistent with recently reported mesoscale tuning organization in macaque body areas (Vanduffel et al., 2023). Finally, qualitative differences in stimulus preference were evident among the two groups of units: neurons aligned with EBA tended to prefer images of human bodies in action (e.g., sports), whereas FBA-aligned neurons preferred animal bodies that included visible faces (Fig. 4J). Together, these findings point to the presence of parallel, functionally distinct sub-populations within macaque body patches, just as face patches (Park et al., 2022).

By aligning macaque electrophysiological responses with human fMRI data under shared naturalistic stimulation, we reveal both cross-species similarities and fine-grained within-region diversity in high-level visual processing. Together, these findings highlight the power of cross-species neuron-voxel alignment for revealing shared representational structures as well as local functional diversity, advancing our understanding of the principles underlying high-level vision in primates.

### Benchmarking Computational Models with Cross-Species Neural Data

The most exciting application of this dataset lies in its potential to bridge neuroscience and artificial intelligence by serving as a large-scale benchmark for evaluating computational models of vision. As a demonstration, we tested the dataset using two classes of models: visual feature models and large language models (LLMs), following approaches used in the human NSD dataset (Doerig et al., 2022) and in recent cross-species studies (Conwell et al., 2025). To systematically assess how well two types of models align with neural representations, we compared their coding spaces using both encoding and decoding analyses.

We began by constructing encoding models using two types of features: visual features, represented by embeddings from the fc6 layer of an ImageNet-pretrained AlexNet, and semantic features, derived from sentence embeddings generated by the MPNet-base-v2 sentence-transformer model (Fig. 5A). To predict neuronal responses, we constructed linear models with cross-validation and evaluated model performance by computing the correlation between predicted and observed responses. Our results indicated that most units were better explained by AlexNet compared to MPNet (example units in Fig. 5B and populational result in Fig. 5C). To quantify relative model performance, we computed the language-visual ratio (LVR), defined as the slope of linear model between predictive performance of the language model and the visual model. We found that LVR of our dataset was 0.74, significantly smaller than 1, indicating that neural responses were more strongly aligned with visual rather than semantic features. However, in the human NSD data, AlexNet and MPNet exhibited similar predictive power, with LVR values close to 1 (Fig. 5D). This suggests a divergence in the relative contributions of visual and semantic representations between species, highlighting a greater integration of semantic information in human high-level visual processing compared to macaques.

**Fig. 5.**
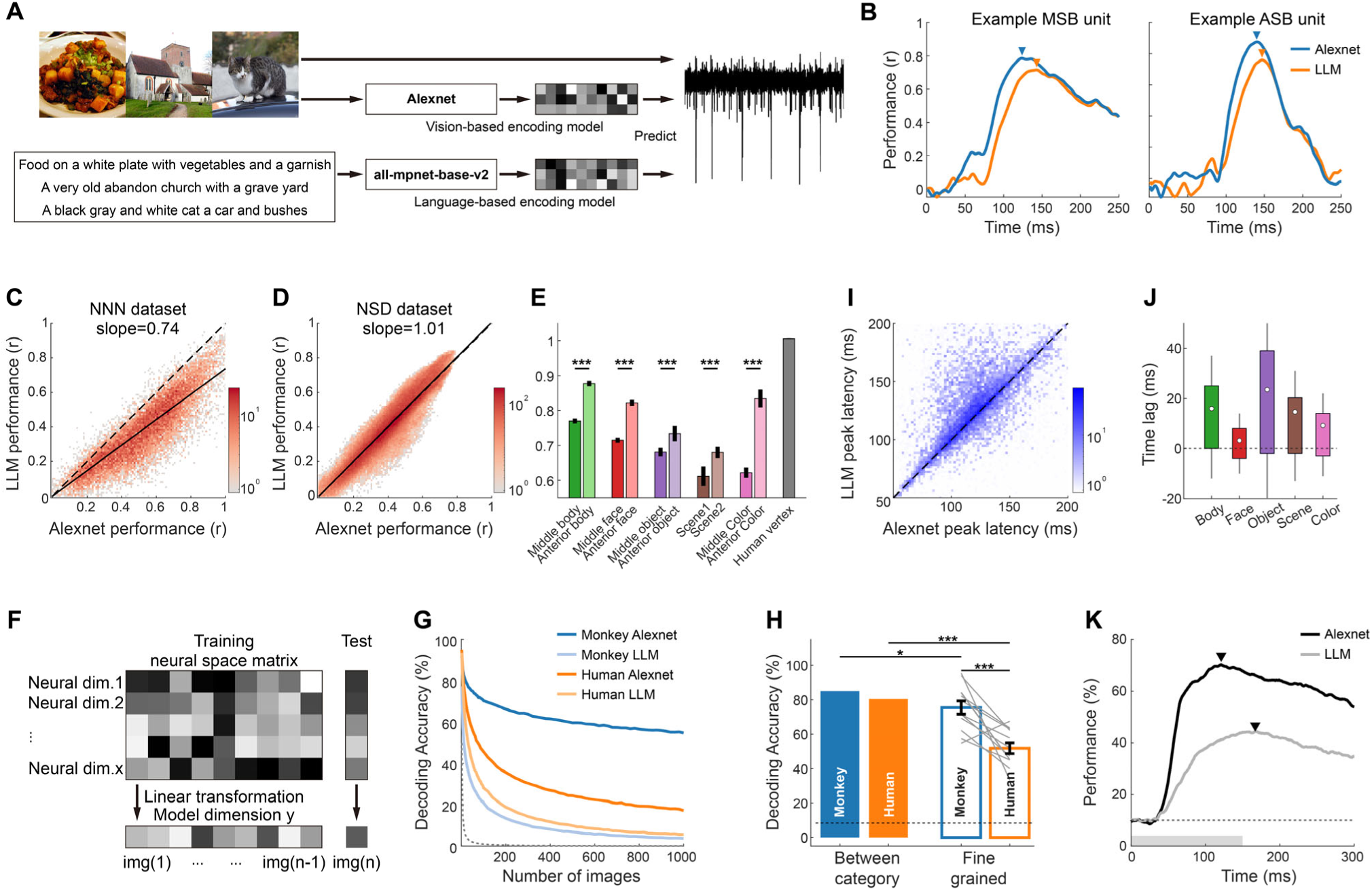
Comparison of coding spaces across species using encoding and decoding analyses. **(A)** Schematic illustration of the neural encoding model. Features were extracted from two sources: the visual model AlexNet, which processes pixel-level image inputs, and the language model MPNet, which encodes semantic captions. These features were used to predict neural responses recorded in the Triple-N dataset. **(B)** Time course of encoding performance for two example units from two body areas. **(C)** Scatter density plot comparing encoding performance on the Triple-N dataset between the visual and LLM models. Each dot represents the log-scale density of unit distribution. The dashed line indicates identical encoding performance across models, while the solid line shows the linear fit. **(D)** Same as **(C)**, but for NSD fMRI dataset. **(E)** Bar plot showing the slope between the two models for each region and the NSD fMRI dataset. Error bars represent three times the standard error of the estimation. **(F)** Schematic illustration of the decoding model. Neural responses were used to predict model features using leave-one-out cross-validation. **(G)** Decoding performance as a function of candidate number across two datasets (Triple-N and NSD) and two coding spaces (Visual and LLM). Dotted lines represent chance performance. **(H)** Twelve-way decoding was performed to match the number of semantic clusters. The left panel shows general decoding performance across clusters, where candidate images were selected from different semantic clusters. The right panel shows within-cluster decoding performance, where candidate images were selected from the same semantic cluster. Gray lines indicate decoding performance for individual clusters; error bars represent the SEM across clusters. Dotted lines denote chance-level performance. **(I)** Scatter density plots of peak latency in encoding performance for both models at log-scale. **(J)** Boxplot of peak latency differences between the two models, where a positive value indicates an earlier peak for the Visual model relative to the language model. **(K)** Decoding performance over time for the two models. Triangles indicate peak time points, with the visual model peaking 47ms earlier than the language model. The dotted line represents chance performance.

Interestingly, when computing the LVR across areas, we observed that LVR values were higher in anterior IT regions relative to middle IT regions across different category-selective networks (Fig. 5E; p<10^-157^ for body areas, p=10^-113^ for face areas, p=10^-9^ for object areas, p=10^-9^ for scene areas, and p=10^-61^ for color areas). This suggests that anterior IT areas may integrate more complex semantic information, whereas middle IT regions remain more strongly tuned to visual features.

Beyond encoding analysis, decoding analysis provides deeper insight into the structure of neural coding spaces by assessing how well stimulus information can be extracted from neural activity (Bao et al., 2020; Chang & Tsao, 2017). We applied decoding to both our dataset and the NSD fMRI dataset: using leave-one-out cross-validation, we trained linear regression models to predict model features from neural responses and decode image identity from the predicted features (Fig. 5F; see **Methods**). Using embeddings from Alenxet model, we achieved significantly higher decoding performance with macaque data than with human fMRI data (Fig. 5G, dark lines). We hypothesize this advantage reflects the finer-grained coding of within-category information at the single-neuron level, which is not accessible with fMRI recordings (Badwal et al., 2025; Dubois et al., 2015). To test this, we evaluated decoding performance within specific semantic clusters (Fig. 5H, see **Methods**). As expected, the decoding performance within the semantic cluster dropped for both species (*t*(11)=2.49, p=0.03 for macaque and *t*(11)=9.1, p<10^-5^ for human) but the decrease was more pronounced in the human NSD data relative to macaque (*t*(11)=4.47, p<10^-3^). This analysis highlights the strength of our dataset in capturing fine-scaled coding of stimuli details at the single-neuron level.

When comparing decoding performance across model spaces, we observed a striking difference divergence. Macaque electrophysiological data achieved higher decoding accuracy with visual features compared to human fMRI data, but lower performance is for semantic features (Fig. 5G, light lines). This dissociation between visual and semantic decoding across datasets is consistent with the LVR findings from encoding analyses. Further, model-dimension-level decoding revealed that human fMRI consistently outperformed macaque data for semantic features, whereas the opposite held true for visual features (Fig. S7A–C). These differences suggest that the coding space in our macaque dataset is more biased toward visual features, whereas the human NSD dataset appears more biased toward semantic features.

Recent studies suggest that neurons in the IT cortex multiplex their functions time windows as brief as 20 ms (Shi et al., 2023), prompting a dynamic comparison of model alignment over time. For encoding analysis, the time to reach peak performance for the language model was significantly later than for the visual feature model (populational results in Fig. 5I and area-wise result in Fig. 5J, p<10^-154^ for body areas, p<10^-10^ for face areas, p<10^-58^ for object areas, p<10^-25^ for scene areas, and p<10^-8^ for color areas). Consistent with this, in the decoding analysis, we found that the visual space peaks earlier than the semantic space across time windows (Fig. 5K, 47ms peak lag for 10 images, with similar trends observed other parameters in Fig. S7E-G). These findings highlight our dataset’s ability to capture the temporally multiplexed functional nature of IT populations, revealing a hierarchical processing sequence where visual features are extracted first, followed by the gradual integration of semantic information.

In summary, the Triple-N dataset provides a powerful benchmark for evaluating computational models of vision, uniquely combining high-resolution single-neuron recordings with naturalistic stimuli. When combined with the NSD dataset, its scale, richness, and cross-species alignment make it ideally suited for testing how well models capture both the spatial and temporal dynamics of neural representations. Together, these datasets position Triple-N as a valuable resource for advancing biologically grounded artificial intelligence.

## Discussion

Our analysis demonstrated the power of fMRI-guided Neuropixels recordings in addressing fundamental questions about object recognition at both population and single-unit levels. By targeting category-selective and non-selective regions in macaque IT cortex (Fig. 1), we uncovered a rich diversity of neural response patterns, revealing distinct temporal dynamics, representational structures, and cross-species similarities and dissimilarities. Within category-selective regions, we confirmed a high concentration of neurons tuned to specific visual categories, supporting the idea that object representations in IT cortex are structured (Fig. 2). Beyond categorical tuning, we identified distinct temporal response types, showing that neuronal activity unfolds over time in a structured manner (Fig. 3). Cross-species comparisons with human fMRI data revealed strong representational alignment between macaque IT and human (VTC). While population-level representations were broadly similar across species, single-unit analyses uncovered substantially greater diversity within macaque IT, including functionally distinct subpopulations with varying tuning profiles and cortical distributions (Fig. 4). Notably, although the core structure of category representations was preserved, we observed species-specific differences in the integration of semantic information: human responses aligned more closely with language-derived features, whereas macaque responses remained more strongly tied to visual feature representations (Fig. 5). Decoding analyses further highlighted the superior granularity of electrophysiological recordings, with macaque neural responses supporting significantly higher decoding accuracy compared to human fMRI (Fig. 5). This underscores the advantage of single-unit resolution in capturing fine-scale stimulus information, particularly for within-category distinctions. Together, these findings demonstrate the value of combining fMRI and Neuropixels recordings to study object recognition, providing Triple-N dataset as a high-resolution, large-scale dataset for understanding neural coding, cross-species vision, and biologically inspired AI models.

FMRI-guided Neuropixels recordings provide us a unique opportunity to tackle the object recognition problem using big data. While fMRI offers a broad, population-level view of category-selective regions, its limited spatial and temporal resolution makes it difficult to decode the fine-scale neural mechanisms underlying object representation (Badwal et al., 2025; Dubois et al., 2015). In contrast, Neuropixels probes enable high-density, single-unit recordings, capturing detailed neuronal activity within these functionally defined regions, however, the spatial coverage of each probe is constrained (∼4 mm along the cortical depth), providing fine-grained resolution within a limited recording volume. By combining the large-scale mapping of fMRI with the fine-grained resolution of Neuropixels, we were able to systematically target and record from a broad set of functionally defined regions across the macaque IT cortex. This approach enabled high-density single-unit recordings within over 20 areas, including both category-selective and non-selective regions, allowing us to capture the detailed temporal dynamics and representational diversity of neuronal populations at scale. Together, our multimodal strategy offers a unique window into the organization of high-level visual processing across spatial and functional hierarchies.

In addition, Triple-N complements the human NSD dataset by offering a high-resolution electrophysiological counterpart to fMRI-based population measurements. While NSD has enabled major advances in image reconstruction and encoding model development using voxel-level data (Takagi & Nishimoto, 2022), its spatial and temporal resolution limits access to fine-scale population dynamics. In contrast, Triple-N dataset captures rich, multi-dimensional neural responses at the single-unit level, enabling more detailed analyses of neural coding, representational geometry, and hierarchical feature processing. This level of resolution opens new possibilities for studying the neural basis of visual perception, including real-time decoding, within-category discrimination, and dynamic population-level computations that are difficult to resolve in fMRI.

Beyond advancing our understanding of biological vision, the Triple-N dataset provides a powerful foundation for developing more biologically inspired neural network architectures. Unlike standard deep neural networks, which are predominantly feedforward and static, the primate visual system relies heavily on recurrent and feedback connections to support dynamic, context-sensitive perception (Kar et al., 2019; She et al., 2024; Shi et al., 2023). Triple-N captures single-neuron activity with high temporal resolution across diverse IT regions, enabling researchers to probe how neural responses unfold over time and how they reflect integrative computations -such as recurrent modulation, or feedback modulations - that are largely absent in current DNNs. By analyzing temporally structured population responses, researchers can identify neural motifs that suggest recurrent loops, top-down modulation, or hierarchical feedforward. Furthermore, because Triple-N aligns with human fMRI data via shared stimuli, it provides a rare opportunity to assess whether such biologically grounded models generalize across species and recording modalities.

Beyond its scientific contributions, this dataset also provides possibility of methodological advancements for neuroscience research. The large-scale, high-density Neuropixels recordings offer a valuable testbed for improving spike-sorting techniques, particularly for refining algorithms that is specialized for primates. Given the depth and breadth of the dataset, it can serve as a benchmark for developing more robust and automated spike-sorting pipelines, advancing data quality in large-scale electrophysiology.

While this dataset provides unprecedented insights into visual representation, several areas could be further improved to enhance its scope and applicability. The rich temporal resolution and single-unit precision of our electrophysiological recordings allow for a fine-grained analysis of neural coding, yet certain aspects of visual processing may require broader stimulus sets or expanded brain coverage to fully capture the complexity of object recognition. One key area for improvement is the stimulus set. Although the Shared1000 images enable direct cross-species comparisons, the number of images remains limited, and the dataset is still biased toward human-centric content. Incorporating a broader range of naturalistic and ecologically relevant stimuli, including dynamic video sequences (Lahner et al., 2024), could provide a more comprehensive view of how visual representations evolve overtime. Additionally, brain coverage could be expanded. While we targeted IT cortex as a key region for high-level visual processing, a more complete understanding of visual coding would benefit from including early visual cortices (Papale et al., 2025) and subcortical areas (dLGN, SC, amygdala). These regions likely play a role in shaping IT representations, and recording from them could offer deeper insights into the hierarchical flow of information in the visual system. By addressing these limitations, future studies could build upon this dataset to develop an even more complete framework for understanding how the brain encodes and processes complex visual information.

In summary, the Triple-N dataset is a valuable and significant resource for the field of visual neuroscience. When combined with other datasets, it has the potential to form a robust ecosystem that will deepen our understanding of visual processing across different scales and species. This synergy will ultimately advance both theoretical models and practical applications in the field, driving new insights into the complex neural networks that underlie vision.

## Methods

### Animals

All experimental protocols followed the guidelines of Institutional Animal Care and Use Committee (IACUC) of Peking University Laboratory Animal Center, and were approved by the Peking University Animal Care and Use Committee. Five male rhesus macaques (Macaca mulatta), aged 5 to 7 years and weighing 5 - 10kg, were used in this study. They were trained to sit in a comfortable, fMRI-compatible chair prior to implant surgery, and were trained to maintain fixation to receive juice rewards during water-control periods. Each subject underwent several separate surgeries under anesthesia. The first surgery involved the implantation of ceramic skull screws (Thomas Recording) and a customized MRI-compatible headpost made from PEEK material. The second surgery was for the implantation of a recording chamber, with the screws, headpost, and recording chamber securely fixed to the skull using dental cement. The recording chamber, made from PEEK material, measures 62 mm from left to right and 50 mm from anterior to posterior (Fig. 1A, coronal view), providing flexibility in probe trajectories while minimizing both the number and size of the craniotomy. The third surgery involved drilling the craniotomy for electrode insertion.

### Behavior paradigm

Stimulus presentation, eye movement recording, and reward delivery were controlled using the NIMH MonkeyLogic (Hwang et al., 2019) toolbox. Eye movements were monitored with infrared eye trackers (JSMZ during the electrophysiology experiment and iSCAN during the fMRI scans). The monkeys were trained to maintain fixation on a small fixation dot to receive a juice reward, with a reward interval of 1 to 4 seconds.

During functional neuroimaging, we employed a block design, with each trial consisting of a 500-ms stimulus ON and 500-ms stimulus OFF period, and a total of 24 trials per block. After each block and before the first block, a 24-second blank block with only a fixation dot was presented. Each functional run included 8 stimulus blocks and 9 blank blocks, lasting a total of 408 seconds and 204 TRs. Stimuli were displayed to the subjects via a video projector with a 60 Hz refresh rate and a spatial resolution of 1024 × 768, positioned 100-120cm from the monkey’s eyes.

For the basic functional localizer, the stimuli comprised 4 blocks, each containing 24 images of faces, bodies, objects, and scrambled images, with a stimulus diameter of 8°. For the scene areas localizer, 4 additional blocks of 96 scene images were included, with a horizontal stimulus size of 18°. For the color areas localizer, 2 blocks of colored, texturized images and 2 blocks of corresponding grayscale images were used, each with a stimulus diameter of 8°.

During the electrophysiology experiment, a small fixation dot was continuously presented at the center of a gray screen (ASUS VG248QE, 24-inch, 1920×1080 resolution, 120 Hz refresh rate) at a distance of 52-54 cm from the monkey’s eyes. Stimuli were presented once the monkey maintained fixation for at least 300 ms. The images followed a 150 ms ON / 150 ms OFF paradigm, with stimulus onset times recorded using a photodiode placed at the corner of the screen. The stimuli consisted of 1000 shared NSD images and 72 localizer images (faces, bodies, objects) from the fMRI localizer, with image diameters ranging from 10° to 12°.

### Magnetic resonance imaging acquisition and processing

The fMRI experiments were conducted using a 3T Siemens Prisma MRI scanner at the Center for MRI Research at Peking University. A customized 8-channel surface coil was used to acquire the MRI data. Prior to the functional scans, a T1-weighted high-resolution 3D structural image was acquired for each subject during an anesthesia session using a 3D-MPRAGE sequence, with TR = 2300 ms, TR = 3.8 ms, flip angle: 9°, number of slices = 224, FOV = 128 mm × 128 mm, voxel size = 0.5 mm × 0.5 mm × 0.5 mm. We used an echo-planar imaging (EPI) sequence for whole-brain functional scans, with TR = 2000ms, TE = 20 ms, flip angle = 80°, number of slices = 27, FOV = 96 mm × 96 mm, voxel size = 1.5 mm × 1.5 mm × 2 mm. To improve the signal-to-noise ratio, monocrystalline iron oxide nanoparticles (MION; Molday ION, 0.26 ml/kg, BioPAL) were injected into the femoral vein of the monkeys prior to the functional experiment.

High-resolution structural images were preprocessed with AFNI’s @animal_warper to nonlinearly registered to NMT v2 template (Jung et al., 2021). Functional scans were preprocessed with AFNI’s afni_proc.py (Jung et al., 2021) or combination of FSL with the Freesurfer Functional Analysis Stream (Fischl, 2012). In short, the preprocessing included field map correction, motion correction and normalization. Only time points during which the monkey maintained fixation were included for design matrix generation. Functional activation was computed using a general linear model, and category selectivity was assessed by performing t-tests contrasting each category against the others.

Functional activation maps, along with structural images from the scanning day or the probe-trajectory structural images (prior to electrophysiology recording, see next section), were co-registered to high-resolution structural images using tkregister2 or align_epi_anat.py for visualization. The inferred recording locations in the macaques’ native space were manually labeled based on the probe-trajectory structural images in 3DSlicer and then registered to a common NMT template using 3dNwarpApply, as shown in Fig. 1B.

### Electrophysiology recording

We designed the probe trajectory based on registered functional activation and high-resolution structural images to ensure precise targeting while avoiding blood vessels. Once the trajectory was finalized, we exported the coordinates relative to the chamber chamber and 3D printed the recording grid for probe placement (Form3, 4, Formlabs). The grid consisted of a base that fit in the chamber and an interface for connecting to the probe. It was also utilized during the craniotomy surgery. Before recording, the printed grid in the chamber and a tungsten probe housed within a plastic guide tube was inserted through the designated grid hole. The recording chamber was then filled with a gadolinium solution for a structural MRI scan to verify the probe location (Fig. 1A).

We recorded neuronal responses from the targeted area using a Neuropixels NHP-Long probe (Trautmann et al., 2023), which is 45 mm long and has 4,416 recording contacts, allowing for 383 simultaneous recording channels. We designed a probe insertion system to ensure stable recordings. In brief, the system included a customized dovetail probe holder mounted on a linear rail, along with an interface that integrated the rail, an oil-hydraulic micromanipulator (Narishige), and components for mounting onto the printed grid. The position of the holder on the rail was controlled by the micromanipulator via a connecting screw. Prior to recording, the probe was installed into a customized probe holder, and the GND and REF channels were shorted using silver wires. The sampling rate for the probe headstage was measured and calibrated before the first recording.

On the recording day, we first inserted a stainless-steel guide tube to penetrate the dura and lowered the probe through the guide tube to the target area. We estimated the probe depth based on the corresponding structural MRI scan and reduced the advancement speed as we were about to approach the targeted location (typically a sulcus or white matter, where the RMS is low; see also Fig.1D for illustration). Once inside the target area, the speed was further reduced to 2-10 μm/s. Typically we waited 20-40 minutes for stabilization of probe. During experiment, we recorded the action potential (AP) and local field potential (LFP) signals using SpikeGLX (billkarsh.github.io/SpikeGLX). Synchronization signals, event codes from the behavioral computer, and photodiode signals were collected using a PXIe-6341 card (NI) along with the electrophysiological signal. For the first 52 sessions, a silver reference line, soldered to a bronze rod and connected to the reference line of the probe, was placed in a saline bath within recording chamber. For the subsequent sessions, we directly connected the probe reference line to the probe holder, which held the guide tube and served as the reference.

### Data analysis: electrophysiological data preprocessing

Electrophysiology data was saved in .bin format using SpikeGLX, behavior data were saved in .bhv2 format using MonkeyLogic. The preprocessing pipeline was carried out using SpikeInterface (Buccino et al., 2020) and custom MATLAB codes. For spike signal, the preprocessing included 300 Hz high-pass filtering, bad channel removal, sampling phase correction, and global median common reference. The processed data were saved for input into Kilosort4 (Pachitariu et al., 2024). We performed spike sorting using Kilosort4 with drift correction, applying a spike detection threshold of 9 for the universal template and 7 for the learned template. Spike quality of sorted units was assessed automatically using BombCell (Fabre et al., 2023), which classified the units as noise, single units (SU), multiple units (MU) and non-somatic units. Non-somatic units were further categorized as SU or MU based on quality features. For LFP signals, preprocessing included bandpass filtering from 1–300 Hz, bad channel removal, sampling time correction, and resampling to 500 Hz.

We only included trials in which the monkeys maintained fixation during the 150ms stimulus presentation period. Spike times from each unit and the processed LFP signals were aligned to each stimulus onset to generate raster plots (Fig. 1E) and LFP plots (Fig. S4C). To generate peristimulus time histograms (PSTH) for each unit in response to the images, we used a 20ms box sliding time window to compute the spike rates.

To assess visual responsiveness, we calculated firing rates for each unit during the baseline period (-25 to 30ms after stimulus onset) and two post-stimulus periods (50-120ms or 120-240ms after stimulus onset). Significance was determined using a two-sided Wilcoxon rank-sum test. Only units with a p-value < 0.001 in at least one post-stimulus periods were included in further analyses, yielding over 30,000 visual responsive units.

### Data analysis: reliability, category selectivity and SNRmax

Due to the variability in response timing across recorded units, we did not use a fixed time window to compute reliability. Instead, we employed a 10 ms sliding window approach, searching over possible start times from 20 to 200 ms and end times from 90 to 390 ms post-stimulus onset, to identify the optimal time window for each unit’s response. With each time window, trials were randomly split into halves and the average response of each half is correlated to estimate the reliability of this units. To cross-validate our method, we used random 500 images as train images to compute the best time window and the left 500 images as test images to validate the reliability, thus comparing the reliability of this method and 50-220ms fixed time window (Fig. S1H). Finally, we repeated this procedure with all 1000 images 10 times and averaged the correlation value r, and applying the Spearman-Brown correction to get the final reliability as:

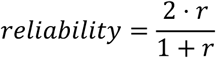

Here r is the Pearson correlation of all images across trials, only units with a corrected reliability greater than 0.4 were included in subsequent analyses, yielding 21,021 reliable responsive units.

We measured categorical selectivity using 72 localizer images. For example, a unit’s face selectivity is calculated as:

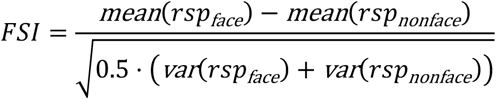

where *rsp*_face_ represents responses to 24 face images, and *rsp*_nonface_ represents these responses to 24 body images and 24 object images (Fig. 2A).

We also computed population-level selectivity to better define the boundaries of categorical selective areas. Using a 200 µm bin, we summed the spike numbers from all units within each local area and then calculated the selectivity (Fig. 2B).

In addition to 72 localizer images, we grouped the 1,000 shared NSD images into 12 semantic clusters using k-means clustering on language model-derived embeddings (see the last section and Fig. S2).

To account for the fact that some neurons exhibited inhibitory responses to non-preferred stimuli (see Fig. 3, Fig. S5), we used the SNRmax method (Papale et al., 2025), to estimate the signal-to-noise ratio. For each unit, the most preferred image was identified based on peak response. SNRmax was then calculated as:

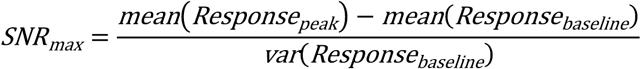

The baseline was defined as the response in a fixed window from −25 ms to +25 ms relative to image onset.

### Response Dynamics

To examine the general response dynamics of each unit, we extracted the PSTH from 0 to 300 ms after stimulus onset and normalized each unit’s time course to have zero mean and unit standard deviation. We then performed k-means clustering to group units based on their temporal response profiles. The silhouette score was used to determine the optimal number of clusters and 3 is the optimal number.

For each cluster, we assessed the temporal coding similarity using representational similarity analysis (RSA). Specifically, for each pair of time points, we computed RDM vectors for the cluster population responses and correlated these vectors to quantify coding similarity across time. In addition, we compared the similarity in responses across different unit types, as shown in Figure S4A.

To examine the spatial organization of PSTH-defined cluster types, we calculated mutual information between cluster identity and the vertical position of each unit within a given area, using the following equation:

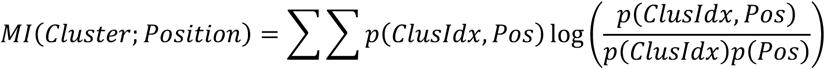

Here, ClusIdx refers to the cluster index i, and Pos is the position of the unit along the probe shank. To evaluate the significance, we permutated the position and re-calculate the mutual information 100 times, the used the median of the permuted MI values as a baseline, A Wilcoxon rank-sum test was then used to compare the observed MI to the permutation distribution.

To further characterize temporal dynamics, we focused on Type 2 units from the anterior face-selective area of monkey M3 as an example. For each stimulus, we computed the mean response and peak response latency, excluding images with normalized average responses below 0.7 to eliminate non-preferred stimuli that may evoke suppressed activity (see Fig. S5). Each unit’s PSTH was normalized by its maximum firing rate before averaging across the population to derive the population response time course.

To evaluate whether peak latency was predictable from image content, we extracted features from AlexNet and reduced them to the first five principal components (PCs). Using leave-one-out cross-validation, we trained a linear regression model to predict each image’s peak latency based on its reduced feature representation. The predicted latencies were then correlated with the observed latencies to assess prediction accuracy.

### The joint analysis for NSD fMRI dataset and Triple-N dataset

We used the NSD dataset and Triple-N dataset to examine cross-species representational similarity. Four human subjects (1, 2, 5, and 7) were selected for analysis, as they completed all experimental sessions. We used the preprocessed, surface-based responses from the ‘betas_fithrf_GLMdenoise_RR’ version on each subject’s native surface, averaging across the three repeated trials per image.

For representational similarity analysis (RSA), we defined the high-level visual cortex mask by taking the union of the ‘nsdgeneral’ ROIs and manually adding category-selective ROIs (EBA, FBA, OFA, FFA, PPA, OPA, OWFA, VWFA) when parts of them were not included in the original NSD general mask. Early visual cortex areas were excluded using the ‘stream’ masks. A representational dissimilarity matrix (RDM) was then computed for each subject within this high-level mask.

In Fig. 5, for the encoding analysis, we selected vertices from the EBA, FBA, OFA, FFA, PPA, and OPA regions in both hemispheres. For the decoding analysis, we included all vertices within the high-level visual cortex mask.

To generate whole-brain correlation maps, we computed the Pearson correlation between macaque neural responses and human voxel responses to the 1,000 shared images, and visualized the results on each subject’s flattened cortical surface, along with manually labeled category-selective regions.

In Fig. 4, to better understand coarse correspondence across species, we included only the top 66% of units whose response profiles most closely resembled the local population. These units were then averaged and treated as representative of the local neuronal response.

To examine the fine-scale diversity of units within category-selective regions, we extracted BOLD responses from the left EBA and left FBA, and computed their partial correlations with the responses of single units in the two body-selective areas of monkey M3. In Fig. S5C, the first 500 images were used to compute the relative difference between the two human brain regions, while Pearson correlation coefficients (PCCs) were calculated on the remaining 500 images to evaluate test-set correspondence.

A unit was classified as an EBA-like unit if it showed a significant (p < 0.01) positive PCC with the EBA response and a non-significant PCC with the FBA response; and vice versa for FBA-like units. To identify preferred images in the body areas, we pooled responses across all body-selective neurons within each region and selected images that elicited a normalized average response greater than 0.5. These are marked as red dots in Fig. 4H.

### Encoding Analysis

For each of the 1,000 shared NSD images, we extracted visual features from the fc6 layer of an ImageNet-pretrained AlexNet (Krizhevsky et al., 2017) model. For semantic features, we used the ‘all-mpnet-base-v2’ model from sentence-transformers, which maps text into a 768-dimensional embedding space, as used in previous studies (Doerig et al., 2022). We generated text embeddings for each of the 5 captions associated with every image and average them to derive semantic feature. The language model was also used to generate semantic clusters by performing k-means clustering on a 768-dimensional space, grouping 1,000 images into 12 labels. These clusters were used for visualization in Fig. 2E and fine-grained decoding in Fig. 5H. The cluster distribution is visualized using t-SNE in Fig. S2.

We constructed encoding models for time-averaged responses using partial least squares (PLS) regression. For each unit, we identified the optimal number of components by minimizing mean squared error (MSE), using a 10-fold cross-validation procedure. Specifically, 900 images were used for training and 100 for testing. Encoding accuracy was quantified as the correlation between predicted and actual responses, normalized by the unit’s response reliability.

To compare the relative bias toward vision versus language models, we fit a linear model *y*=*βx* using MATLAB’s lscov function, where *x* represents the performance of the AlexNet encoding model and *y* the MPNet encoding model. The slope (language-vision ratio) was compared across brain regions within the same network (e.g. face network) using t-tests. To analyze the temporal dynamics of the encoding models, we reduced the model dimensionality to 60 principal components with PCA and built linear encoding models for the response profiles at each time point using lscov function.

### Decoding Analysis

To compare the decoding performance, the following neural responses were selected: (1) time-averaged population response from all reliable macaque units, (2) Surface-based beta values from the high-level visual cortex of all human NSD subjects (3) time-resolve responses from all reliable macaque units. For fair comparison, the response spaces were first reduced to 500 dimensions (response dimension), and the visual or semantic features were reduced to 100 dimensions (model dimension) with PCA. We then performed leave-one-out cross-validation: for each of the 1,000 images, we trained a linear regression model using the response dimensions from the other 999 images to predict model feature dimensions, and used the fitted weights to estimate the model features for the held-out image.

To quantify the decoding performance of each dimension, the Pearson correlation between predicted and actual dimension to 1000 images were calculated. To assess image-wise decoding accuracy, we randomly selected a number of candidate images and compared the Euclidean distance between predicted and actual feature vectors. A decoding trial was counted as correct if the predicted vector was closest to its true counterpart. This process was repeated 50,000 times to estimate accuracy. To evaluate the ability of each neural representation to support between-cluster and within-cluster discrimination, we performed decoding using candidate images drawn either from 12 different semantic clusters or from the same cluster, using the same procedure.

## Data and code availability

Preprocessed trial-wise time course data (in .h5 format), unit-related data such as spike waveform and reliability, raw behavioral data (in .bhv2 format), and raw neural data (∼4TB, in .bin format), and analysis code will be made available to the corresponding author during the peer review process and on a public platform upon publication.

## Supplementary Figures

**Fig. S1.**
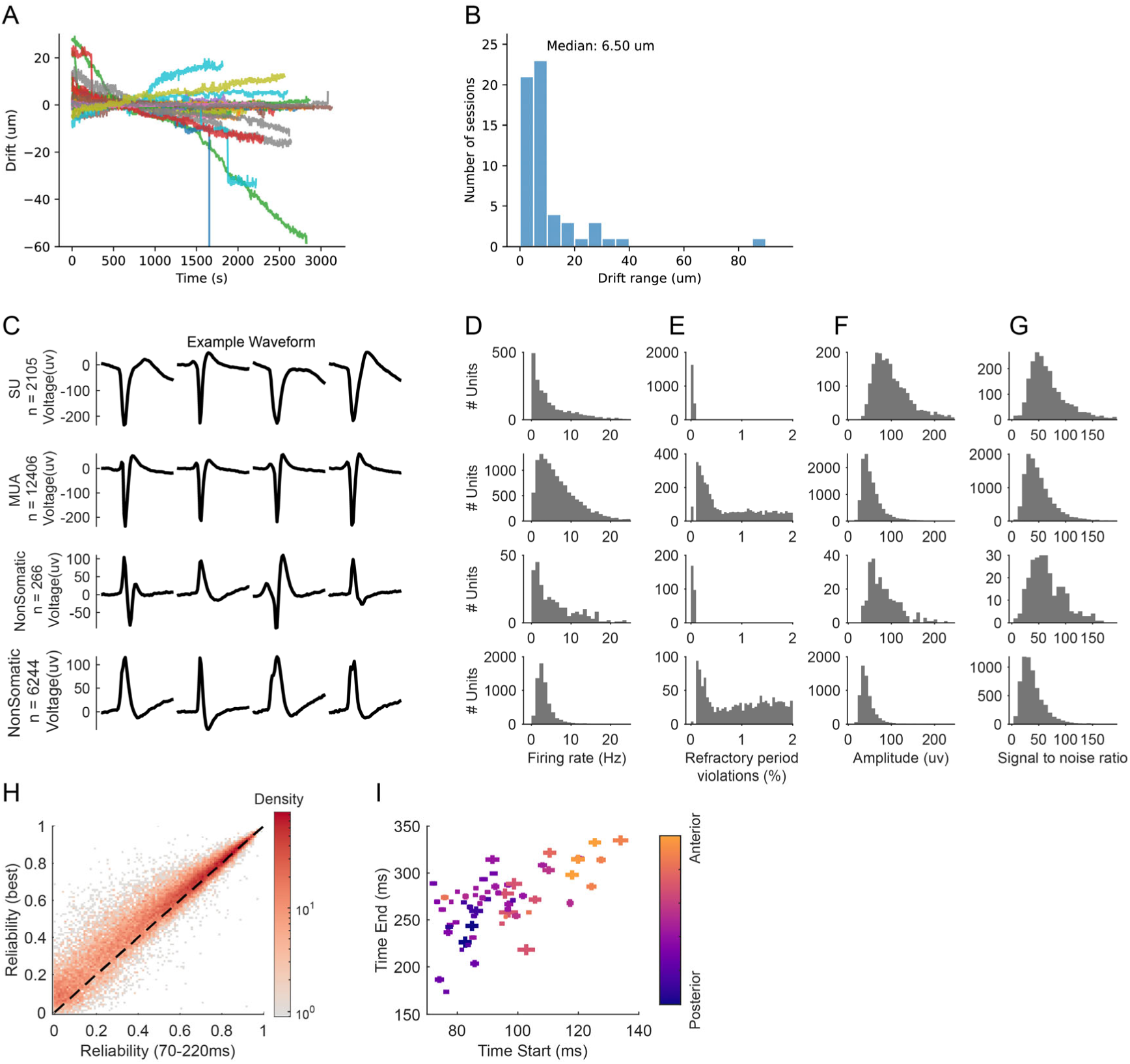
Quality metrics across the entire dataset. **(A)** Estimated probe drift over experiment time, with each line representing a single session. **(B)** Distribution of probe drift range across all sessions. **(C–G)** Spike quality metrics, from top to bottom: single-unit activity, multi-unit activity, non-somatic single-unit, and non-somatic multi-unit activity. **(C)** Four Example spike waveforms for each group. **(D)** Firing rate over the course of the session. **(E)** Percentage of refractory period violations. **(F)** Spike waveform amplitude. **(G)** Spike waveform signal-to-noise ratio. **(H)** Scatter plot of response reliability (Spearman-Brown corrected correlation) for fixed time windows (x-axis) versus searched time windows (y-axis), with density represented in color on log10 scale. **(I)** Scatter plot of searched time window across areas, with color indicating the anteroposterior coordinate of the NMT template. Error bars represent the SEM across units.

**Fig. S2.**
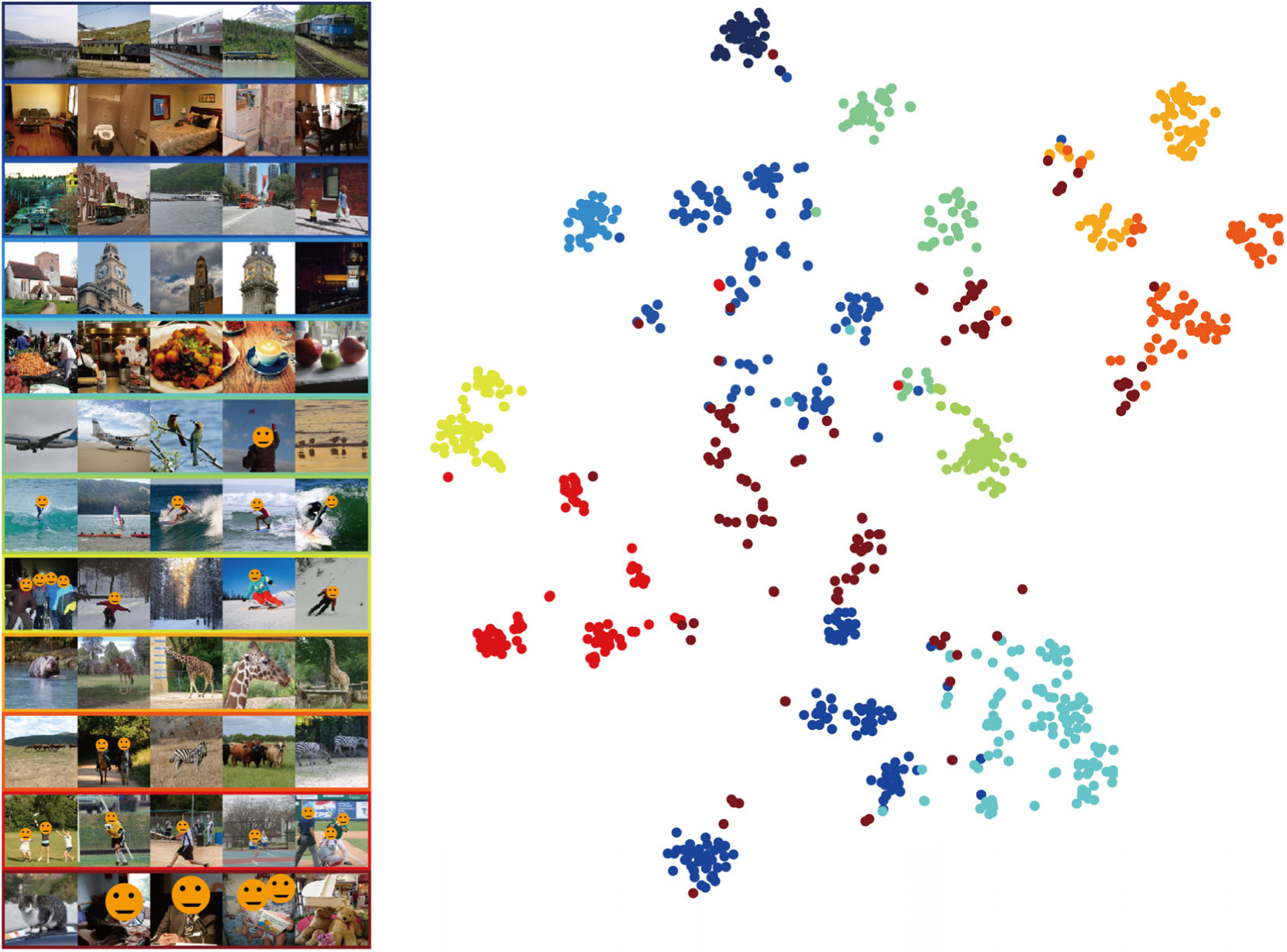
Semantic clusters of 1000 images from LLM embedding. The cluster is used for visualization in Fig.2 and fine-grained decoding in Fig. 5. Due to the bioRxiv policy, all face images were obscured with a mask.

**Fig. S3.**
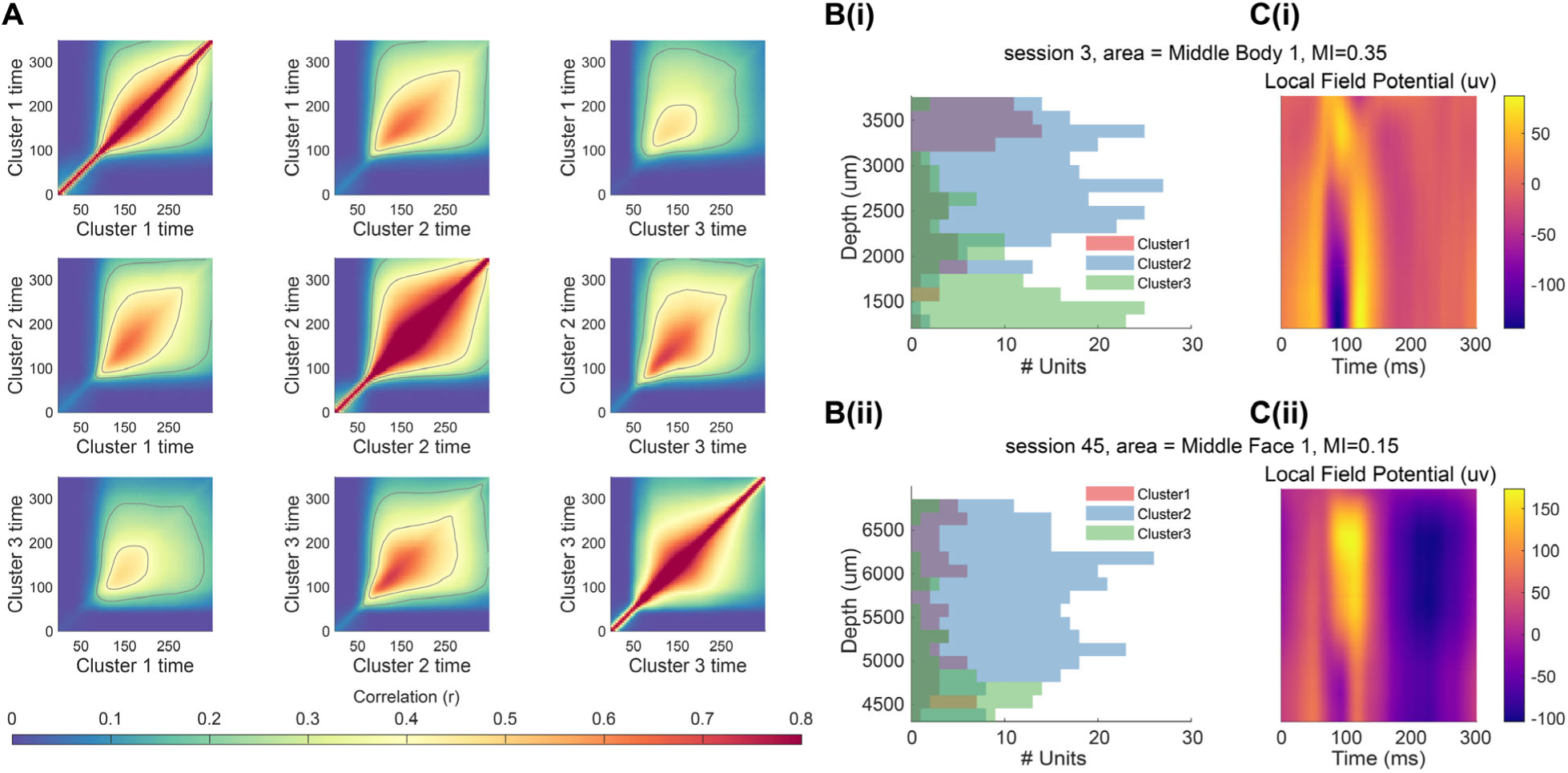
Temporal representational structure and anatomical distribution of neuronal response types. **(A)** Cross-population representation similarity across time. Each matrix represents the similarity between three unit types in Fig.3B. The grey lines indicate contour levels at 0.2 and 0.4. **(B-C)** Two example sessions illustrating the spatial distribution of unit types, with mutual information values between unit type and position as 0.35 (upper example) and 0.15 (lower example). **(B)** Histogram showing the distribution of unit types as a function of cortical depth. **(C)** LFP signals across shank positions.

**Fig. S4.**
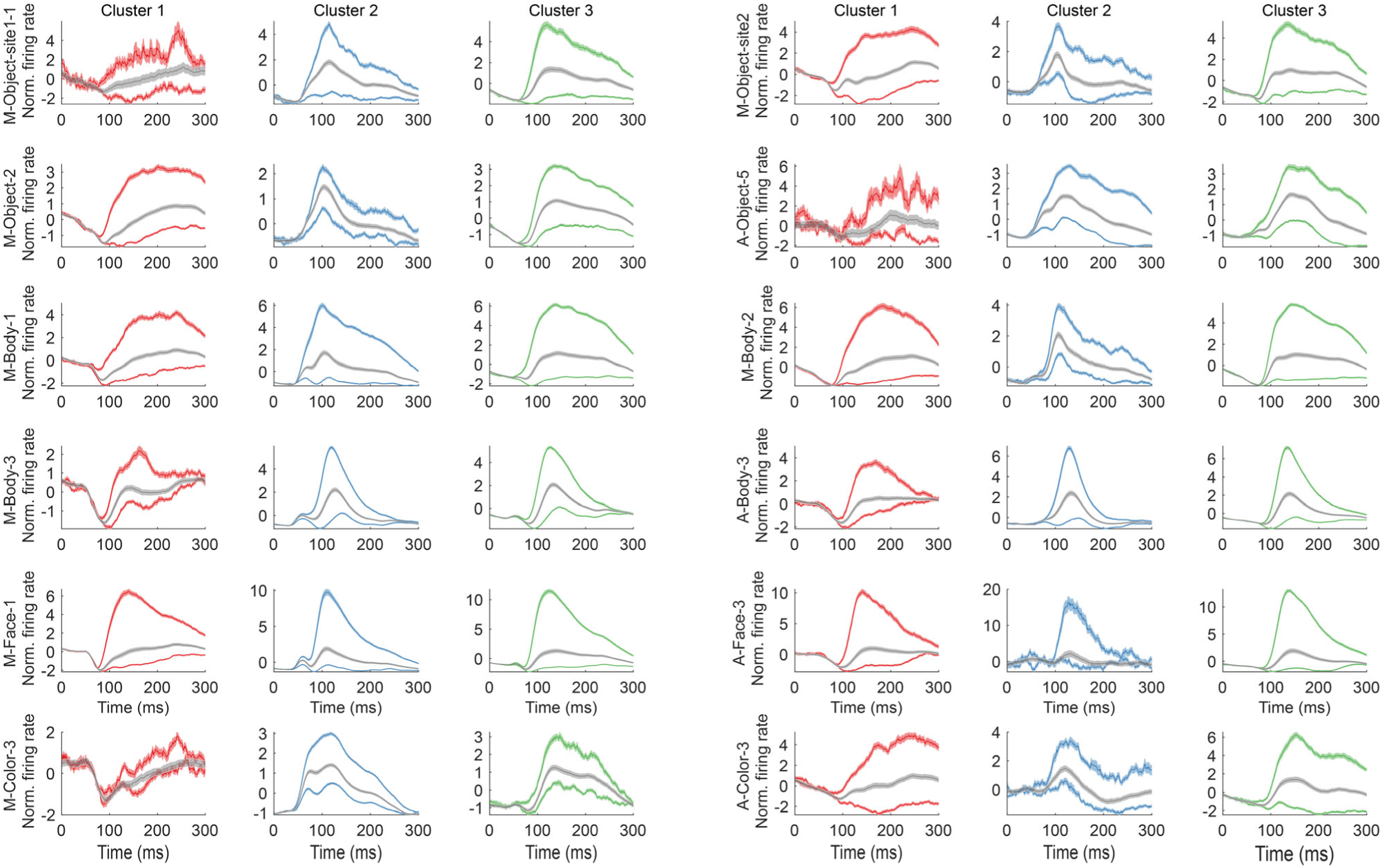
Example response profiles from three types of units. The response profiles are shown for: (1) 50 most preferred images, indicated by colored lines above the grey lines; (2) the averaged response, represented by the grey lines; and (3) 50 least preferred images, indicated by colored lines below the grey lines, for each area.

**Fig. S5.**
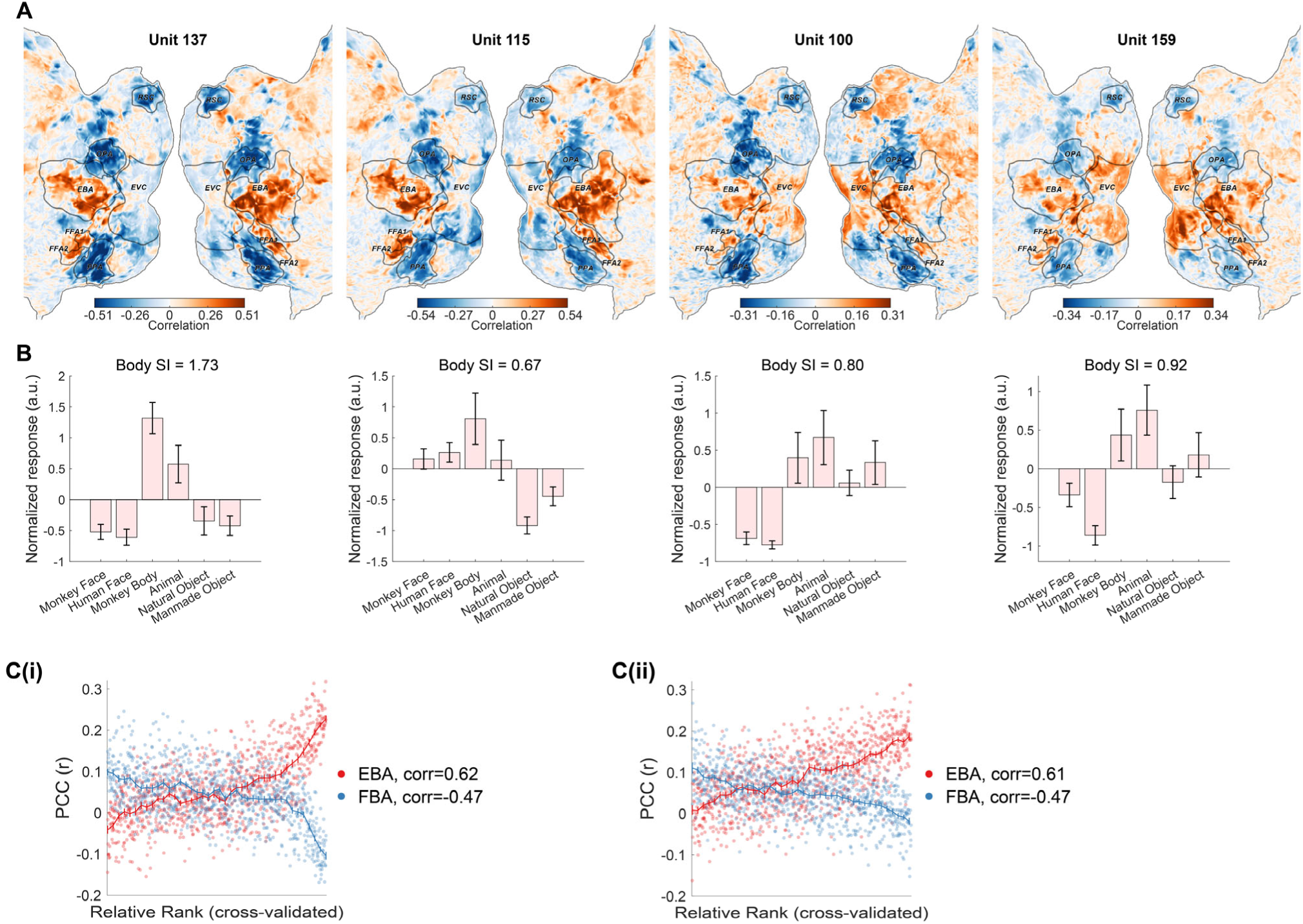
Within-region diversity revealed by cross-species comparison. **(A)** Whole-brain correlation map of four example units with NSD subject 5, normalized by response reliability. **(B)** Response profiles of the four example units to the localizer stimuli, along with their body selectivity. Error bars indicate SEM across 12 images. **(C)** Partial correlation between unit responses and human ROIs as a function of cross-validated rank. Dots represent individual units, while lines indicate the median from 200-unit bins, **(Ci)** for the middle body patch, **(Cii)** for the anterior body patch.

**Fig. S6.**
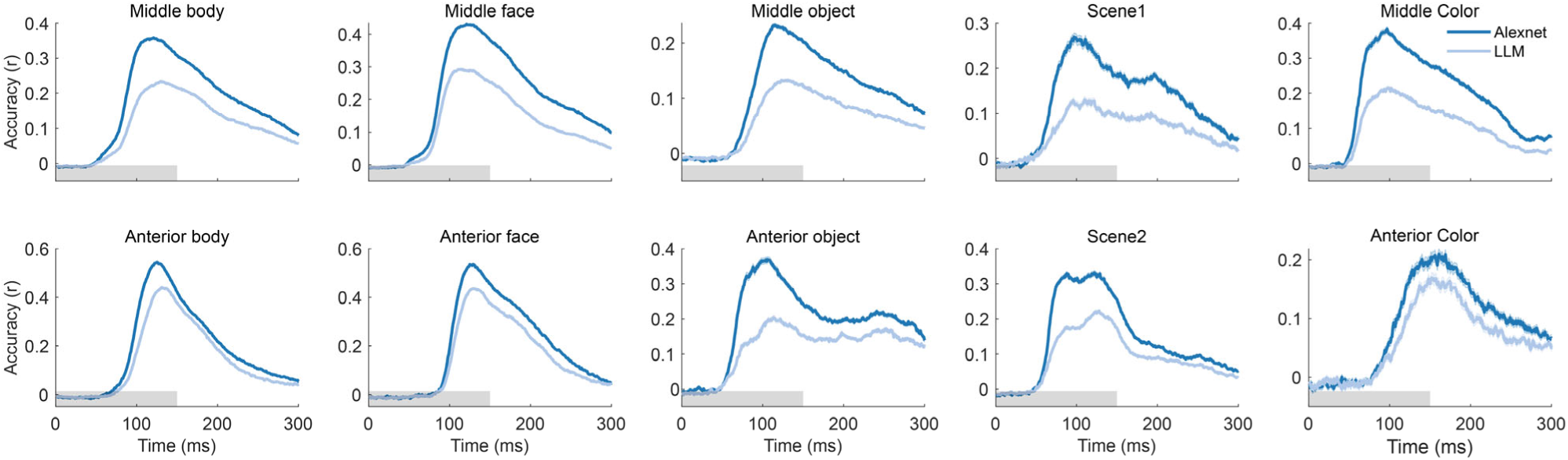
Temporal profiles of encoding performance for the visual and language models in each recorded area.

**Fig. S7.**
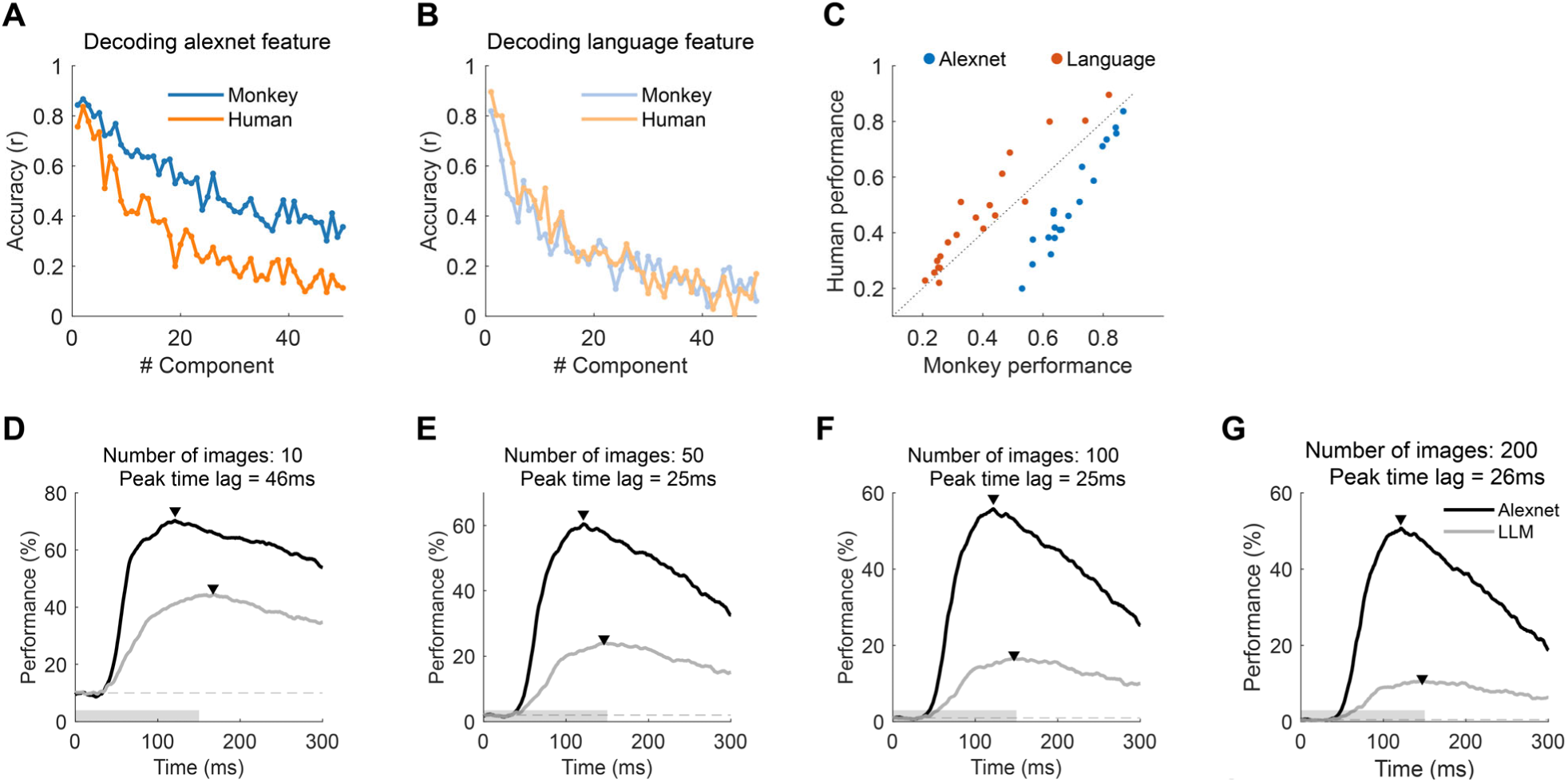
Decoding analysis across model space. **(A)** Decoding performance of feature representations, using principal components from the visual model, evaluated across both the macaque electrophysiology and human fMRI datasets. **(B)** Same as **(A)**, but for principal components from the language model. **(C)** Scatter plot of decoding performance (correlation) for the first 20 principal components across two datasets, with the grey line indicating identical performance. **(D-G)** Decoding performance across time for various number of candidate images. Triangles indicate peak time points, and dotted lines represent chance performance.

